# Insulin signaling accelerates the anterograde movement of Rab4 vesicles in axons through Klp98A/KIF16B recruitment via Vps34-PI3Kinase

**DOI:** 10.1101/2024.04.24.590898

**Authors:** Kamaldeep Singh, Semanti Das, Sabyasachi Sutradhar, Jonathon Howard, Krishanu Ray

**Affiliations:** Department of Biological Sciences, Tata Institute of Fundamental Research, Mumbai - 400005, India; Department of Molecular Biophysics and Biochemistry, Yale University, New Haven, CT - 06520, United States; National Brain Research Centre, Manesar, Haryana – 122051, India

**Keywords:** Rab4 vesicles, insulin signaling, phosphoinositide-3-phosphate [PI(3)P], Klp98A, Vps34, axonal transport, synaptic homeostasis, neurodegeneration, kinesin, recycling endosomes

## Abstract

Rab4 GTPase organizes endosomal sorting essential for maintaining the balance between recycling and degradative pathways. Rab4 localizes to many cargos whose transport in neurons is critical for regulating neurotransmission and neuronal health. Furthermore, elevated Rab4 levels in the CNS are associated with synaptic atrophy and neurodegeneration in *Drosophila* and humans, respectively. However, how the transport of Rab4-associated vesicles is regulated in neurons remains unknown. Using *in vivo* time-lapse imaging of *Drosophila* larvae, we show that activation of insulin signaling via Dilp2 and dInR increases the anterograde velocity, run length, and flux of Rab4 vesicles in the axons. Molecularly, we show that activation of neuronal insulin signaling further activates Vps34, elevates the levels of PI(3)P on Rab4-associated vesicles, recruits Klp98A (a PI(3)P-binding kinesin-3 motor) and activates their anterograde transport. Together, these observations delineate the role of insulin signaling in regulating axonal transport and synaptic homeostasis.

**Highlights:**

- Dilp2-mediated insulin signaling activates anterograde transport of Rab4 vesicles
- Vps34 regulates anterograde velocity, run length and flux of Rab4 vesicles
- Local PI(3)P signaling on Rab4 vesicles regulates their motility in the axons
- PI(3)P production upon acute insulin stimulation recruits Klp98A on Rab4 vesicles

## Introduction

Rab4 is a critical organizer of endosomal sorting and plays a crucial role in several processes like recycling of surface receptors (van der Sluijs et al., 1992), axon outgrowth (Falk et al., 2014), maintenance of spine morphology, and synaptic plasticity (Hoogenraad et al., 2010). Rab4 activation induces its recruitment onto vesicles which subsequently activates the vesicle trafficking through kinesin and dynein motors. Rab4 localizes to and transports vesicles carrying various cargos like tetraspanins (Moretto et al., 2023), neuregulins (Ahmad et al., 2022), integrins (Shin et al., 2021), and astrotactins (Behesti et al., 2018) regulating integral neuronal processes like trafficking and degradation of surface proteins, synapse formation, maturation, and maintenance. Unsurprisingly, a few of these cargo are also implicated in neurodevelopmental (Behesti et al., 2018) and psychiatric disorders (Ahmad et al., 2022). On the other hand, overactivation of Rab4 at the presynaptic terminals induces synaptic atrophy in *Drosophila*. In humans as well, brain autopsies of Alzheimer’s patients have indicated that elevated levels of Rab4 in cholinergic basal forebrain (CBF) and CA1 neurons of hippocampus are inversely correlated with their cognitive abilities (Ginsberg et al., 2010, 2011). Thus, the activation of Rab4 must be tightly controlled and must not be too great as to disturb neuronal homeostasis. Therefore, understanding the yet unidentified molecular mechanisms regulating the movement of Rab4-associated vesicles (henceforth called Rab4 vesicles) in neurons is essential for better comprehension of membrane trafficking functions in health and disease.

Besides the widely studied roles of insulin signaling in regulating metabolism (Tokarz et al., 2018), growth (Hakuno & Takahashi, 2018), and development (Dupont & Holzenberger, 2003), a few studies have also shown that insulin regulates intracellular transport in peripheral tissues (Imamura et al., 2003; Kumar et al., 2019). For instance, insulin activates GLUT4 recycling in adipocytes by activation of Rab4 and recruitment of kinesin-2 (Imamura et al., 2003) and the anterograde transport of lipid droplets by phosphatidic acid-dependent recruitment of kinesin-1 in hepatocytes (Kumar et al., 2019). Furthermore, insulin signaling in the central nervous system (CNS) is implicated in APP and HAP1 transport (Shieh et al., 2020) and regulates AMPA receptor endocytosis and long-term depression (LTD) in hippocampal neurons (Man et al., 2000). Insulin signaling in the brain is also associated with cognitive decline and aging in *Drosophila* and humans (Akintola & Van Heemst, 2015; Augustin et al., 2017). We therefore hypothesized that insulin signaling may play a role in regulating the axonal transport of Rab4 vesicles.

Here, using *in vivo* time-lapse imaging coupled with genetic and pharmacological perturbations, we show that Dilp2 and dInR-mediated insulin signaling accelerates the anterograde axonal transport of a subset of Rab4 vesicles in *Drosophila* cholinergic neurons. Further, using a combined RNAi and chemical inhibitor screen of all known Phosphatidylinositol-3-Kinases (PI3Ks) in *Drosophila*, we show that this process is directly mediated by Vps34 (Class-III PI3K). Activation of Vps34 by insulin signaling leads to the production of Phosphatidylinositol-3-Phosphate [PI(3)P] on Rab4 vesicles, which finally recruits Klp98A, a kinesin-3 family motor on Rab4-associated vesicles and activates their anterograde movement. In effect, we elucidated the molecular basis of activating anterograde axonal transport of Rab4 vesicles in neurons by insulin signaling which may play a role in maintenance of synaptic homeostasis.

## Results

### Treatment with human Insulin and Dilp-2 increases the anterograde flux, velocity, and run length of Rab4 vesicles in cholinergic neurons of *Drosophila*

Previous studies have shown that acute insulin stimulation activates Rab4 and increases the surface levels of GLUT4 in a kinesin-2 and PI3K-dependent manner (Imamura et al., 2003; Shibata et al., 1997). In addition, Rab4 activity, PI3K, and kinesin-2 motor are implicated in Rab4 vesicle transport in the axons (Dey et al., 2017). Therefore, we hypothesized that insulin stimulation will activate the anterograde transport of Rab4-associated vesicles in the axons. To test this hypothesis, we chose third instar *Drosophila* larvae as a model system. We have previously shown that both endogenous and fluorophore-tagged Rab4 (Rab4mRFP) exhibit a similar localization and enrichment in the ventral ganglion of third instar *Drosophila* larvae (Dey et al., 2017).

*In vivo* time-lapse imaging of cholinergic neurons expressing Rab4-mRFP revealed that acute stimulation (15 minute) with 1.7 nM human insulin in the bath could visibly increase the movement of Rab4 vesicles in axons (Fig. 1A, Supplemental Movie S1). A detailed estimation revealed a significant increase of anterograde flux by ∼10% and a proportional reduction of the retrograde flux upon insulin stimulation (Fig. 1A and 1B; n≥19 segmental nerves, N≥3 larvae). Amongst eight *Drosophila* insulin-like peptides (Dilps) (Nässel & Broeck, 2016), Dilp2 and Dilp5 are the most abundantly produced insulin-like peptides in the brain (Semaniuk et al., 2021) with Dilp2 having the highest homology to human insulin with 35% sequence identity (Brogiolo et al., 2001). Therefore, to understand the physiological relevance of the effect, we next tested the effect of recombinant *Drosophila* insulin-like peptides 2 and 5 (Dilp2 and Dilp5) on axonal transport of Rab4 vesicles. Like human insulin, acute stimulation with 17 nM Dilp2 also increased the net anterograde flux of Rab4 vesicles (Fig. 1A and 1B; n≥19 segmental nerves, N≥3 larvae). On the contrary, acute stimulation with Dilp5 even at 10-fold higher concentrations had no significant effect on the anterograde flux of Rab4 vesicles, indicating a certain degree of specificity in Dilp2 action (Fig. 1, S1A-C, and Table S1; n≥14 segmental nerves, N≥3 larvae). These observations suggested that both human insulin and Dilp2 stimulations could selectively increase the anterograde flux of Rab4 vesicles in *Drosophila* axons, albeit human insulin was more effective, owing likely to its enhanced stability (*Phoenix Pharmaceuticals*, n.d.; *Sigma*, n.d.).

**Figure 1:**
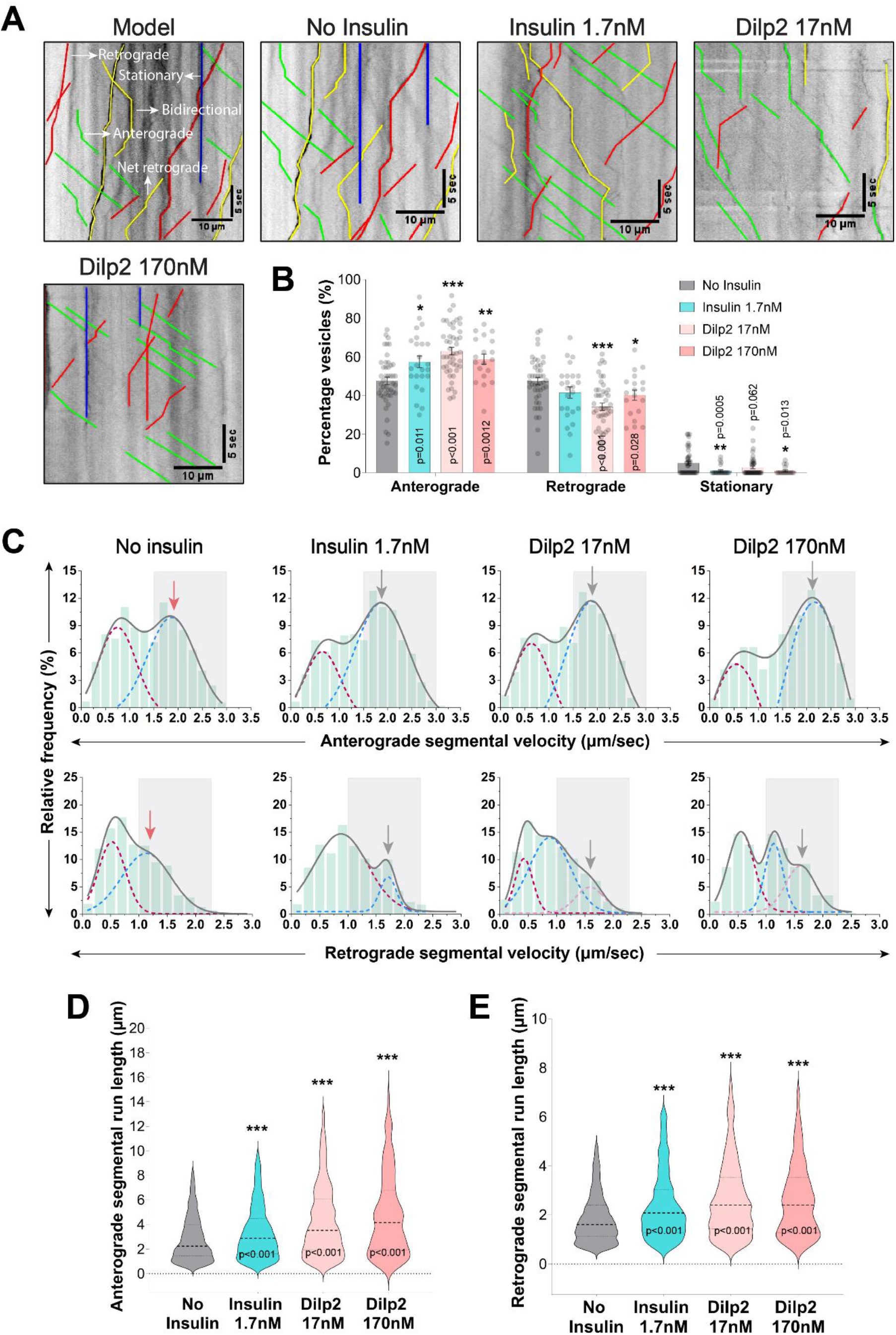
Acute treatment with Human Insulin and Dilp2 increases the anterograde flux, velocity, and run length of Rab4 vesicles in *Drosophila* cholinergic neurons. **A)** Representative kymographs from different treatment groups depicting anterograde (green), retrograde (red), stationary (blue), and bidirectional (yellow) trajectories of Rab4 vesicles obtained from the time-lapse images depicted in Supplemental Movie S1. **B)** Percentage of Rab4 vesicles divided in the net anterograde, retrograde, and stationary groups based on the net displacement (in 1 minute) across different treatment groups. All statistical comparisons are to the ‘No Insulin’ control group and are done using Mann-Whitney U test. **C)** Frequency histograms of anterograde (top row) and retrograde (bottom row) segmental velocities (µm/sec) of Rab4 vesicles across different treatment groups. Grey solid lines indicate the cumulative fit of multimodal distribution while the maroon, blue, and pink dashed lines indicate individual gaussian distributions (see Methods for details). The fast-moving runs have been marked with a gray box in both anterograde (1.5 - 3.0 µm/sec) and retrograde (1.0 - 2.25 µm/sec) velocity plots and the red and grey arrows indicate the peak value in the fast-moving range for the control and test group(s), respectively. Comparisons to the ‘No Insulin’ control group are done using Kolmogorov-Smirnov (KS) test for comparing sample distributions. **D and E)** Violin plots depicting anterograde (D) and retrograde (E) segmental run length (µm) of Rab4 vesicles across different treatment groups. All statistical comparisons are to the ‘No Insulin’ control group and are done using Mann-Whitney U test. (p values are mentioned on the plot only in cases where the comparison is statistically significant).

The increased anterograde flux of Rab4 vesicles could be promoted by, a) enhanced anterograde velocity or processivity, b) inhibition of retrograde movement, or c) combination of both. To identify which of these factors were at play, we analysed the transport parameters of individual segmental runs i.e., the segmental run length and velocities of Rab4 vesicles under different conditions. The Rab4 vesicles moved with a bimodal distribution of anterograde segmental velocities (like what was observed previously for Rab6 vesicles in hippocampal sensory neurons (See methods for details; Gumy et al., 2017) divided arbitrarily into two categories for ease of understanding: 0.0 – 1.5 µm/sec and 1.5 – 3.0 µm/sec, hereafter referred to as slow and fast-moving runs, respectively (Fig. 1C). A similar bimodal distribution was observed in the retrograde direction and was divided into slow (0.0 – 1.0 µm/sec) and fast-moving runs (1.0 – 2.25 µm/sec) (Fig. 1C). Quantification revealed that insulin treatment (both Dilp2 and human insulin) significantly increased the frequency of fast-moving runs in the anterograde direction (Fig. 1C and Table S1; n≥800 runs, N≥3 larvae), and the anterograde segmental run length (Fig. 1D; n≥800 runs, N≥3 larvae). It also increased the segmental run length and velocity in the retrograde direction, although to a lesser extent compared to the anterograde direction (Fig. 1C and E, and Table S1; n≥340 runs, N≥3 larvae). Together, these data show that Dilp2/insulin stimulation preferentially enhances the anterograde motility of the Rab4 vesicles in the axons, thus indicating a surprising link between insulin signaling and vesicle transport in axons.

### Cell-autonomous insulin signaling activates anterograde transport and selectively increases the anterograde flux of Rab4 vesicles

To understand whether the observed effect of insulin on transport of Rab4 vesicles is mediated by cell-autonomous insulin signaling, we specifically knocked down endogenous *Drosophila* insulin receptor (dInR) in the cholinergic neurons using a previously validated RNAi line (Jia et al., 2015). While dInR RNAi significantly reduced the anterograde flux of Rab4 vesicles by ∼8% (Fig. 2A; n≥14 segmental nerves, N≥3 larvae), overexpression of constitutively active insulin receptor (InR^CA^) in the cholinergic neurons significantly increased the anterograde flux of Rab4 vesicles by ∼10% (Fig. 2A; n≥20 segmental nerves, N≥3 larvae). This increase in anterograde flux was abolished by acute treatment with a well-known chemical inhibitor of insulin signaling, LY294002, which inhibits all known PI3Ks inside the cell (Fig. 2A; n≥19 segmental nerves, N≥3 larvae). InR^CA^ overexpression also enhanced the frequency of fast-moving runs in both the anterograde (∼15%; Fig. 2B; n≥600 runs, N≥3 larvae) and retrograde direction (∼10%; Fig. S2A; n≥550 runs, N≥3 larvae), which is consistent with the results obtained with acute stimulation of Dilp2/insulin earlier (Fig. 1A-B). Further, the frequency of fast-moving runs decreased both in the InR RNAi (∼16%; Fig. 2B; n≥750 runs, N≥3 larvae) and LY294002 treatment in the InR^CA^ backgrounds (∼14%; Fig. 2B and Table S1; n≥200 runs, N≥3 larvae). Similarly, the segmental run lengths in the anterograde direction decreased upon InR RNAi and increased in the InR^CA^ backgrounds, respectively (Fig. 2C; n≥200 runs, N≥3 larvae). The increase in anterograde segmental run-lengths was suppressed by the inhibitor treatment (Fig. 2C; n≥750 runs, N≥3 larvae). Consistent with the previous observations (Fig. 1), the effects on the retrograde segmental velocities were not so prominent in the InR^CA^ background and only a small but visible change was observed in the InR RNAi and LY294002 treatment (in the InR^CA^ background) conditions (Fig. S2; n≥200 runs, N≥3 larvae). Together, these observations suggest that activation of cell-autonomous insulin signaling via dInR increases the anterograde speed and processivity of Rab4 vesicles resulting in a significant enhancement of the anterograde flux of these vesicles in the cholinergic neurons of *Drosophila* (Fig. 2D). In contrast, its effect on the retrograde movement of Rab4 vesicles was not conclusive.

**Figure 2:**
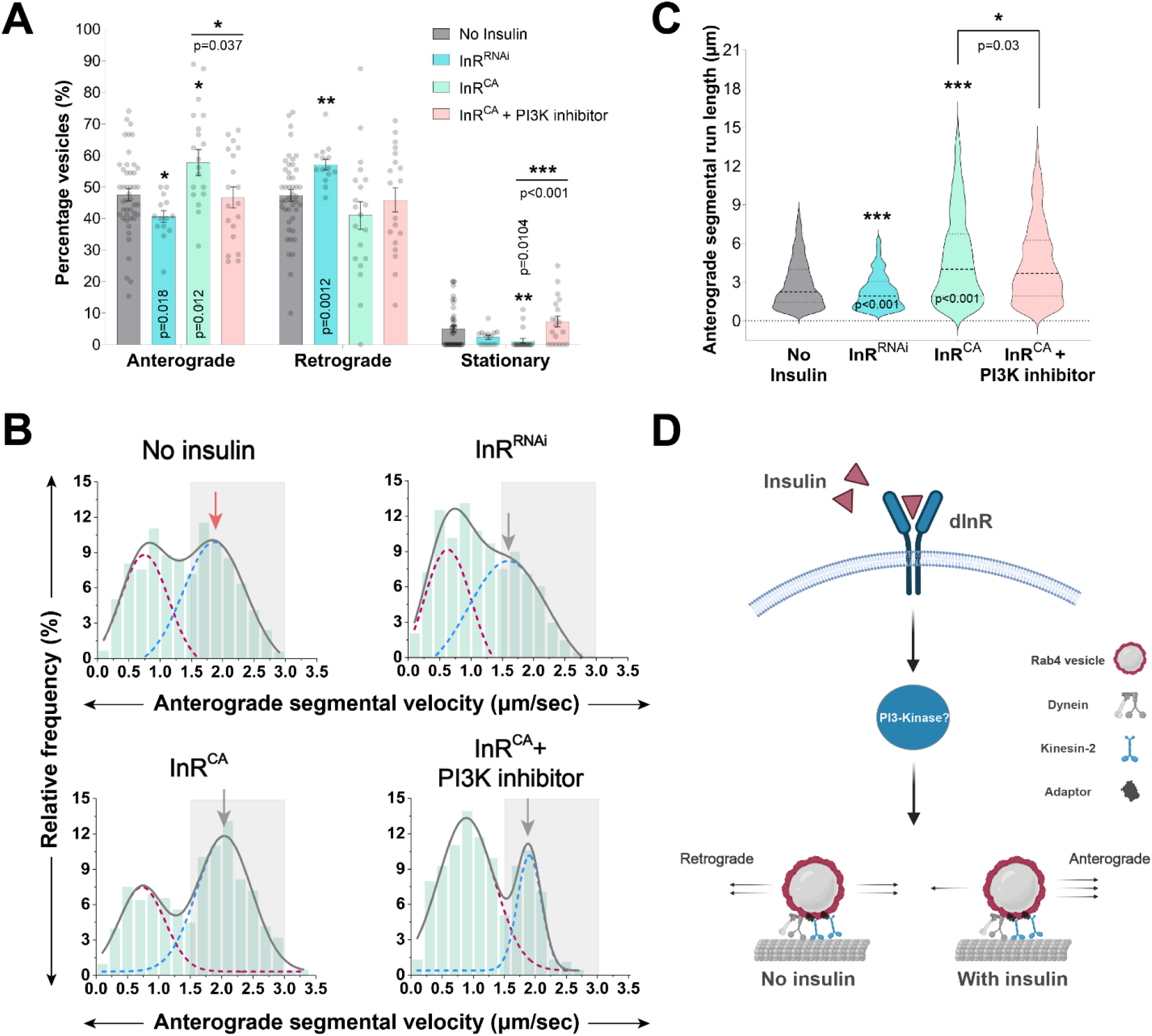
Cell-autonomous insulin signaling regulates anterograde flux, velocity, and run length of Rab4 vesicles through a PI3K-dependent pathway. **A)** Percentage of Rab4 vesicles divided in the net anterograde, retrograde, and stationary groups based on the net displacement (in 1 minute) across different genetic and pharmacological perturbations (See also Supplemental Movie S2 for reference). Each dot indicates flux from one segmental nerve bundle (Collected from 3-7 animals in all experiments henceforth). Note that there is a significant reduction in the anterograde flux in InR^RNAi^ and a significant increase in the anterograde flux in InR^CA^, which is inhibited upon acute treatment with PI3K inhibitor. All statistical comparisons are to the ‘No Insulin’ control group (unless specified otherwise) and are done using the Mann-Whitney U test. **B)** Frequency histograms depicting anterograde segmental velocity (µm/sec) of Rab4 vesicles from different treatment groups. Grey solid lines indicate the cumulative fit of bimodal distribution while the red and blue dashed lines indicate the individual gaussian distributions. The fast-moving anterograde runs (1.5 - 3.0 µm/sec) have been marked with a gray box in the velocity plot and the red and grey arrows indicate the peak value in the fast-moving range for the control and test group(s), respectively. Comparisons to the ‘No Insulin’ control group are done using Kolmogorov-Smirnov (KS) test. **C)** Violin plots depicting anterograde segmental run length (µm) of Rab4 vesicles from different treatment groups. All statistical comparisons are to the ‘No Insulin’ control group unless specified otherwise and are done using the Mann-Whitney U test. **D)** Schematic depicting signaling cascade where Dilp2 via dInR through a PI3K-dependent pathway activates and increases the anterograde movement of Rab4 vesicles in *Drosophila* cholinergic neurons. Dynein motor can bind directly to Rab4(Bielli et al., 2001) and kinesin-2 binds Rab4(Dey et al., 2017) through yet unidentified accessory proteins. (p values are mentioned on the plot only in cases where the comparison is statistically significant.)

### Vps34 activity downstream to insulin signaling regulates anterograde flux, velocity, and run length of Rab4 vesicles

Previous studies have indicated that PI3K regulates Rab4 activity and turnover in adipocytes and neurons, respectively (Dey et al., 2017; Imamura et al., 2003). In addition, reduced anterograde transport upon acute treatment with a pan-PI3K inhibitor in InR^CA^ overexpression-background (Fig. 2A) suggested an active role of phosphoinositide(s)-mediated lipid signaling in regulating Rab4 transport in the axons. Three different classes of PI3Ks are expressed in *Drosophila* sensory neurons (Brown et al., 2014; H. Li et al., 2022) and they localize to distinct organelles inside the cell (Di Paolo & De Camilli, 2006). Therefore, to identify the specific class of PI3K acting downstream to insulin signaling involved in the activation of anterograde transport of Rab4 vesicles, we conducted a combinatorial screen using specific chemical inhibitors and cell-specific knockdown of all the PI3Ks.

Acute inhibition (15 minute) of all classes of PI3Ks using a pan-PI3K inhibitor (LY294002) significantly reduced the anterograde flux (∼14%; Fig. 3A; n≥10 segmental nerves, N≥3 larvae), the frequency of fast-moving anterograde runs (∼22%; Fig. 3B and Table S1; n≥350 runs, N≥3 larvae), and the run length (Fig. 3C) of Rab4 vesicles. Likewise, acute inhibition of Class III PI3K (Vps34) using a specific inhibitor (SAR405) led to a similar effect (Figure 3A-C and Table S1), thereby indicating a possibly direct role of Vps34 in this process. In contrast, acute inhibition of the Class-I PI3K using a specific inhibitor (HS173) did not affect the anterograde flux (Fig. 3A), although it had a significant effect on the frequency of fast-moving runs (Fig. 3B and Table S1). Consistent with the effects of insulin treatments (Fig. 1 and Fig. S2), we noted only small effects on retrograde transport parameters after the inhibitor treatments (Fig. S3). Altogether, these findings suggest that Vps34 could selectively regulate the anterograde flux of Rab4 vesicles downstream of insulin signaling in axons. To confirm these conclusions, we individually knocked-down all three PI3Ks in cholinergic neurons. It revealed that only the Vps34 knockdown could significantly reduce the anterograde flux by nearly 10% (Fig. 3D and S4; n≥17 segmental nerves, N≥3 larvae), as well as the frequency of fast-moving runs (∼11%; Fig. 3E; n≥750 runs, N≥3 larvae) and the run length (Fig. 3F) of the Rab4 vesicles. Furthermore, acute insulin stimulation in the Vps34 knockdown background failed to increase the anterograde flux of Rab4 vesicles to the expected level (Fig. 3D; n≥18 segmental nerves, N≥3 larvae) and only marginally improved the frequency of fast-moving anterograde runs (∼4%; Fig. 3E; n≥850 runs, N≥3 larvae). Lastly, RNAi of the PI3KC1-catalytic subunit had no significant effect on the anterograde flux as well as the frequency of fast-moving anterograde runs and anterograde run length (Fig. S4; n≥20 segmental nerves, N≥3 larvae). However, RNAi of the PI3KC1-regulatory subunit or the PI3K-C2 marginally decreased the frequency of fast-moving anterograde runs, and significantly increase both the anterograde and retrograde run length (Fig. S4; n≥6 segmental nerves, N≥3 larvae), though the effect was much less compared to that of Vps34 knockdown. Taken together, these results suggest that downstream of insulin signaling, Vps34 acts to stimulate the anterograde movement of Rab4 vesicle in axons.

**Figure 3:**
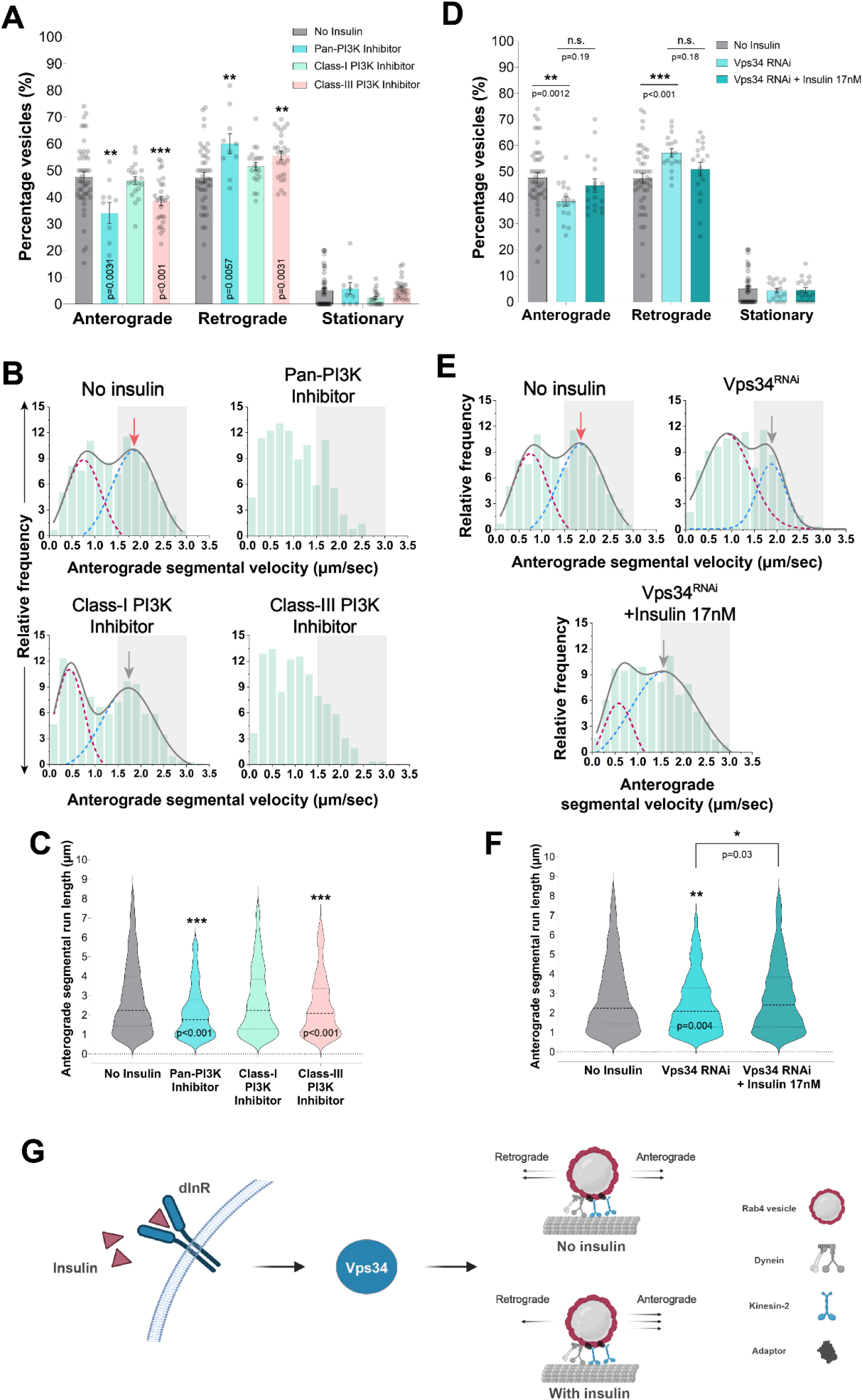
Acute inhibition and knockdown of Class-III PI3K/Vps34 reduces anterograde flux, velocity, and run length of Rab4 vesicles in the axon. **A)** Percentage of Rab4 vesicles divided in the net anterograde, retrograde, and stationary groups based on the net displacement (in 1minute) across different treatment groups (see Supplemental Movie S3). All statistical comparisons are to the ‘No Insulin’ control group and are done using Mann-Whitney U test. **B)** Frequency histograms depicting anterograde segmental velocity (µm/sec) of Rab4 vesicles across different treatment groups. Grey solid lines indicate the cumulative fit of bimodal distribution while the red and blue dashed lines indicate the individual gaussian distributions from the bimodal histogram. Curve fitting has been omitted in certain groups due to lack of an optimal fit and the fast-moving anterograde runs (1.5 - 3.0 µm/sec) have been marked with a gray box in all the velocity histograms. The red and grey arrows indicate the peak value in the fast-moving range for the control and test group(s), respectively. Comparisons to the ‘No Insulin’ control group are done using Kolmogorov-Smirnov (KS) test. **C)** Violin plots depicting anterograde segmental run lengths (µm) of Rab4 vesicles across different treatment groups. All statistical comparisons are to the ‘No Insulin’ control group unless specified otherwise and are done using Mann-Whitney U test. **D)** Percentage of Rab4 vesicles divided in the net anterograde, retrograde, and stationary groups based on the net displacement across different genetic and pharmacological perturbations (see Supplemental Movie S4). **E)** Frequency histograms depicting anterograde segmental velocity (µm/sec) of Rab4 vesicles across different genetic and pharmacological perturbations. **F)** Violin plots depicting anterograde segmental run lengths (µm) of Rab4 vesicles. All statistical comparisons are to the ‘No Insulin’ control group (unless specified otherwise) and are done using Mann-Whitney U test. **G)** Schematic depicting a summary of signaling cascade where Vps34 activation is essential for the overall increase in the anterograde movement of Rab4 vesicles in the axons downstream to insulin stimulation. (p values are mentioned on the plots only in cases where the comparison is statistically significant.)

### Vps34-dependent PI(3)P-mediated lipid signaling regulates velocity of Rab4 vesicles in axons

Vps34 is an early endosome-localized (Sato et al., 2001) lipid kinase that produces PI(3)P from phosphatidylinositol (PtdIns) both *in vitro* (Bago et al., 2014) and *in vivo* (Schu et al., 1993). Further, cellular PI(3)P can be readily detected by a genetically-encoded 2x-FYVE-GFP biosensor (Pattni et al., 2001). Therefore, to identify the presence and Vps34-dependent enhancement of the PI(3)P levels on the mobile Rab4 vesicles, we used simultaneous, dual-colour, time-lapse imaging of Rab4mRFP vesicle movement in cholinergic neurons expressing Rab4-mRFP and 2x-FYVE-GFP. It showed that PI(3)P biosensor marked the motile Rab4mRFP vesicles in the axons in both the anterograde and retrograde directions (Fig. 4A-B and Supplementary Movie S10; N=5). While about 10% of the total vesicles (n=14 ROIs, N=3 larvae, >8000 total vesicles; See Methods for details) were dually marked with Rab4mRFP and 2xFYVE-GFP, acute insulin stimulation increased the percentage of PI(3)P-Rab4 colocalized vesicles by ∼10% (Fig. 4C and 4D; n≥11 ROIs, N=3 larvae, >7000 total vesicles) consistent with our previous findings (Fig. 1B). Importantly, the insulin-dependent PI(3)P increment on Rab4 vesicles was abrogated by an acute treatment with the Vps34-specific inhibitor SAR405 (Fig. 4C and 4D; n≥9 ROIs, N≥3 larvae, >1000 total vesicles). Together, we conclude that activation of Vps34 downstream of neuronal insulin signaling promotes the formation of PI(3)P on Rab4 vesicles, which in turn increases the anterograde movement of Rab4 vesicles.

**Figure 4:**
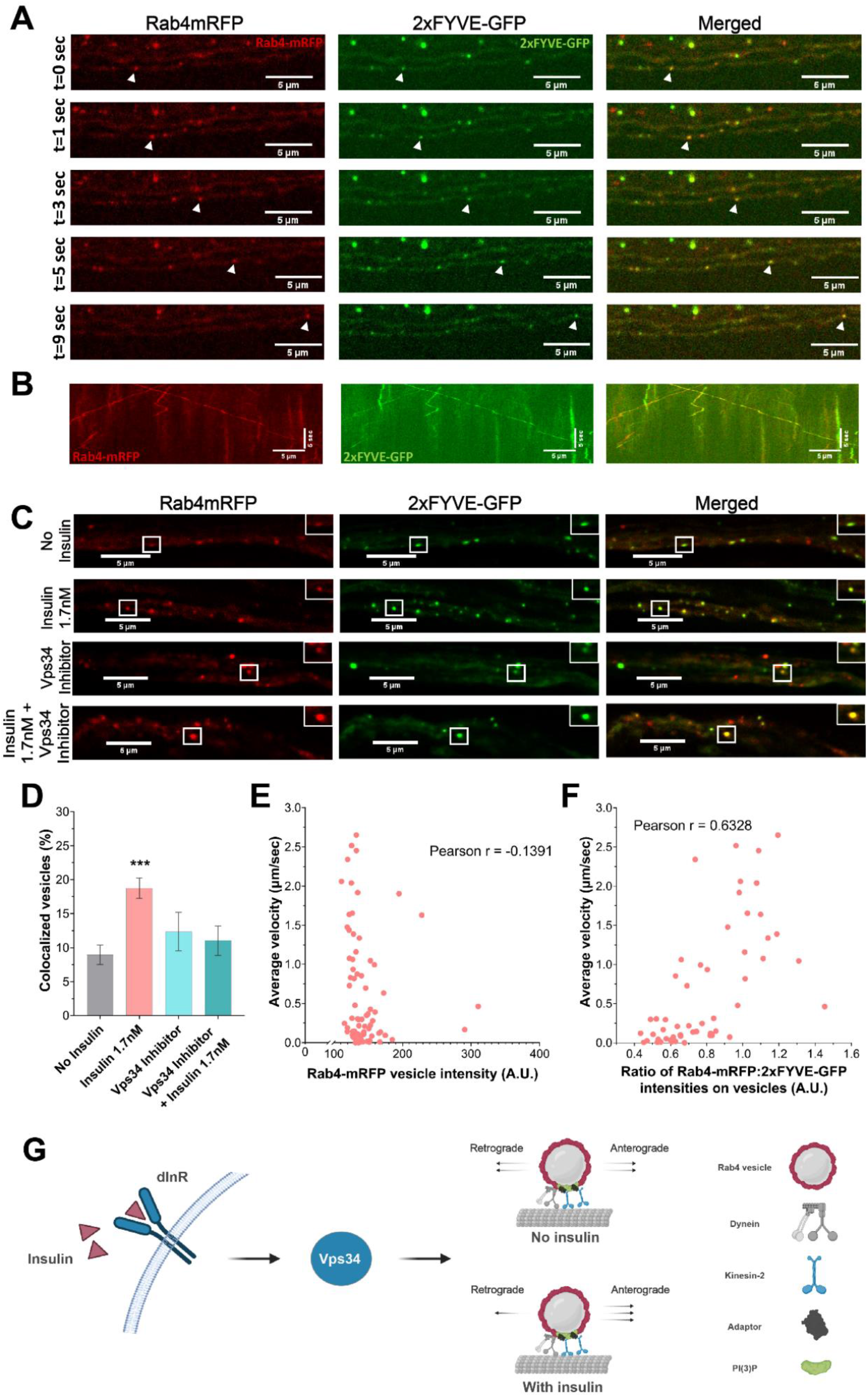
Activation of local PI(3)P-mediated lipid signaling by insulin stimulation regulates transport of Rab4 vesicles. **A)** Time-lapse images of segmental nerve axons depicting comigration of Rab4 (marked in red) and genetically-encoded PI3P biosensor 2xFYVE-GFP (marked in green; see Supplemental Movie S10). **B)** Kymographs illustrates bidirectional movement of Rab4 vesicles (marked in red) with the PI3P biosensor 2xFYVE-GFP (marked in green. **C)** Representative images from larval fillet preparations under different pharmacological perturbations (for 15minutes) post-fixation depicting the levels of Rab4 (marked in red) and PI3P biosensor 2xFYVE-GFP (marked in green in axons. **D)** Bar graph depicting percentage of Rab4-mRFP (marked in red) and 2xFYVE-GFP (marked in green) colocalized vesicles across different pharmacological perturbations. All statistical comparisons are to the ‘No Insulin’ control group and are done using Mann-Whitney U test. Mean ± SEM plotted. **E)** Scatter plot showing no correlation between Rab4-mRFP (marked in red) intensity on individual vesicles with average velocity (µm/sec) of the same vesicles obtained using single-particle tracking. The value of Pearson Correlation Coefficient (r) is mentioned on the plot. **F)** Scatter plot revealing a positive correlation between Ratio of Rab4-mRFP:2xFYVE-GFP on the same vesicles as (E) with average velocity (µm/sec) of the same vesicles obtained using single-particle tracking (see Supplemental Movie S12). The value of Pearson Correlation Coefficient (r) is mentioned on the plot. **G)** Schematic depicting a summary of signaling cascade where insulin-dependent Vps34 activation leads to an increase in PI(3)P production on Rab4 vesicles. Masking of PI(3)P-binding domain by 2xFYVE-GFP biosensor also disrupts overall motility of vesicles indicating that local PI(3)P-mediated lipid signaling on the surface of Rab4 vesicles is essential for the overall increase in the anterograde movement of Rab4 vesicles downstream to insulin stimulation. (p values are mentioned on the plot only in cases where the comparison is statistically significant).

Formation of PI(3)P on the endosome surface activates PI(3)P-mediated lipid signaling (Tsukazaki et al., 1998). Therefore, we wanted to investigate the role of PI(3)P-mediated lipid signaling in regulating the axonal transport of the Rab4 vesicles. For this purpose, we harnessed the documented ability of the PI(3)P biosensor to mask the PI(3)P domain on the surface of endosomes and prevent the binding of endogenous effectors (Gillooly, 2000), thus abrogating PI(3)P-mediated lipid signaling. Therefore, we hypothesized that the levels of 2x-FYVE-GFP on a Rab4 vesicle could proportionately abrogate PI(3)P-signaling and accordingly affect the transport properties of the vesicle if PI(3)P is the causal factor behind the enhanced motility of Rab4 vesicles. A simultaneous estimation 2x-FYVE-GFP and Rab4mRFP intensities on individual vesicles in the axons coupled with single-particle tracking of the same vesicles indicated no correlation between the Rab4mRFP levels on vesicles (a proxy for Rab4 activation) and the average segmental velocities (Fig. 4E; n=76 vesicles, N=6 larvae). In contrast, levels of the PI(3)P biosensor were inversely correlated with the average velocity of these vesicles (Fig. 4F; n=76 vesicles, N=6 larvae) i.e. higher the intensity of biosensor, lower the velocity of the vesicle and vice versa (See Supplemental Movie S12). In other words, abrogation of PI(3)P-mediated lipid signaling on Rab4 vesicles leads to a proportional reduction in their motility. Together with the previous results, this observation further established that local PI(3)P-signaling downstream to activation of insulin signaling on the surface of Rab4 vesicles is a critical determinant of their anterograde transport properties, likely by regulating recruitment of kinesin motors on the vesicles (Fig. 4G).

### Loss of kinesin-2 function failed to block the insulin-mediated increased anterograde transport of Rab4 vesicles

We next asked if heterotrimeric kinesin-2, a key regulator of Rab4 vesicle transport in cholinergic neurons (Dey et al., 2017) and insulin-mediated transport of GLUT4-carriers in adipocytes (Imamura et al., 2003), plays a role in the observed increase in anterograde transport downstream of insulin signaling. Using dual-color live imaging of cholinergic neurons expressing Rab4mRFP and Klp64D-GFP (KIF3A ortholog in *Drosophila* and an essential subunit of kinesin-2), we demonstrate that kinesin-2 is likely to comigrate with Rab4 vesicles in the axons (Fig. 5A-B and Supplemental Movie S9; N=2). However, these events were infrequently detected due to very high cytoplasmic backgrounds of Klp64D-GFP (Fig. 5A).

**Figure 5:**
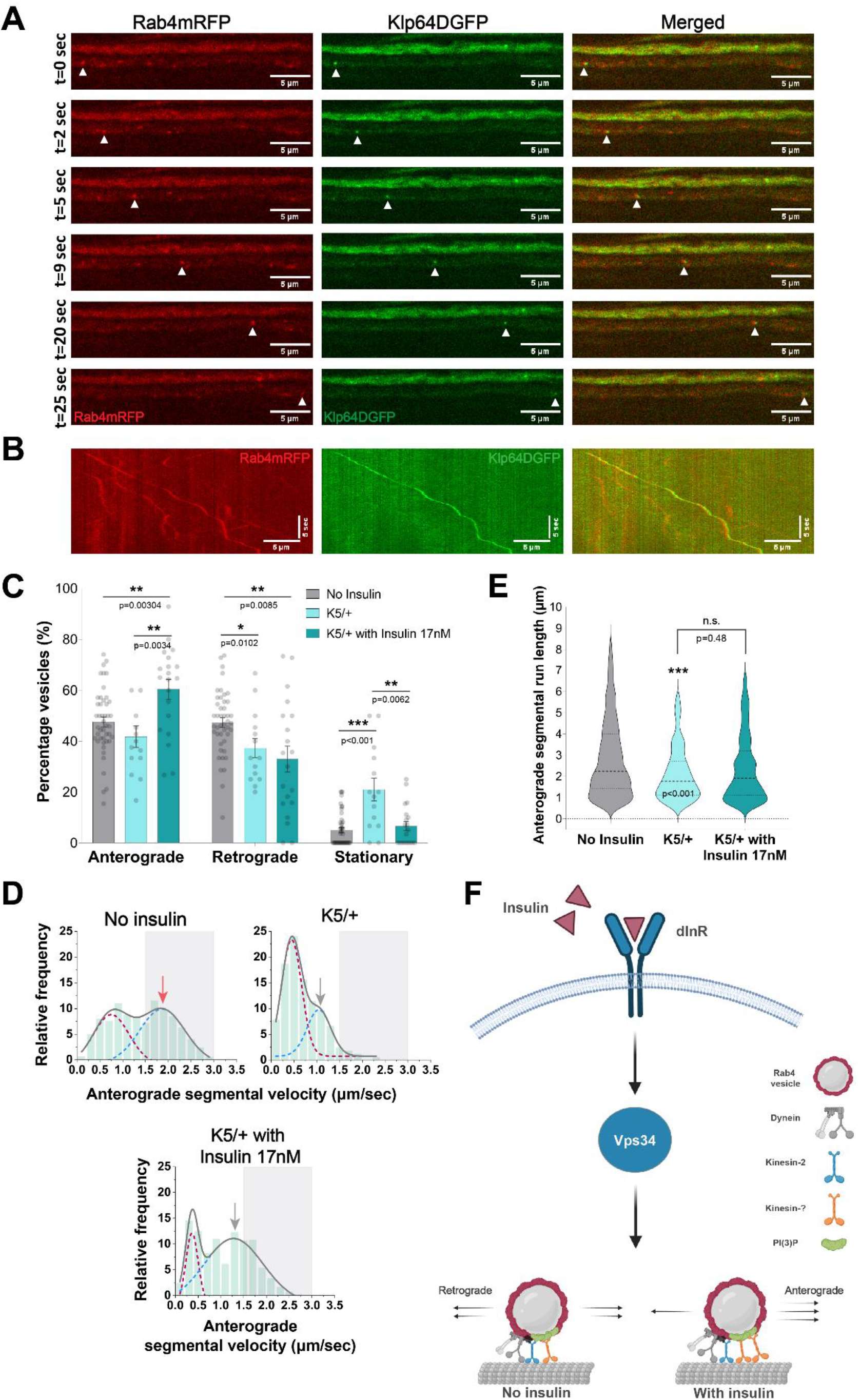
Kinesin-2 does not regulate the increase in anterograde flux of Rab4 vesicles in the axons downstream of insulin signaling. **A)** Time-lapse images of segmental nerve depicting comigration of Rab4 vesicles (marked in red) and Klp64D, KIF3A ortholog (marked in green; see Supplemental Movie S9). **B)** Kymographs illustrates movement of Rab4 vesicles (marked in red) with the Klp64D (marked in green) in segmental nerve axons of cholinergic neurons. **C)** Percentage of Rab4 vesicles divided in the net anterograde, retrograde, and stationary groups based on net total displacement (in 1minute) across different genetic and pharmacological perturbations (see Supplemental Movie S5). All statistical comparisons are to the ‘No Insulin’ control group and are done using Mann-Whitney U test. **D)** Frequency histograms depicting anterograde segmental velocity (µm/sec) of Rab4 vesicles across different treatment groups. Grey solid lines indicate the cumulative fit of bimodal distribution while the red and blue dashed lines indicate the individual gaussian distributions from the bimodal histogram. The fast-moving anterograde runs (1.5 - 3.0 µm/sec) have been marked with a gray box in all the velocity plots and the red and grey arrows indicate the peak value in the fast-moving range for the control and test group(s), respectively. Comparisons to the ‘No Insulin’ control group are done using Kolmogorov-Smirnov (KS) test for comparing sample distributions. See also Table S1. **E)** Violin plots depicting anterograde segmental run lengths (µm) of Rab4 vesicles across different genetic and pharmacological perturbations. All statistical comparisons are to the ‘No Insulin’ control group (unless specified otherwise) and are done using Mann-Whitney U test. **F)** Schematic depicting a summary of signaling cascade where insulin-dependent Vps34 activation leads to increase in the PI(3)P levels on Rab4 vesicles but kinesin-2 does not play a role in the increased anterograde flux of Rab4 vesicles. (p values are mentioned on the plot only in cases where the comparison is statistically significant)

Further, to perturb the function of kinesin-2, we used a previously characterized *Klp64D* mutant (*Klp64D*^*K5*^) known to affect kinesin-2 function *in vivo*(Ray et al., 1999). Consistent with the previous report in lch5 neurons of *Drosophila* which showed a significant decrease in the anterograde flux and segmental velocity of Rab4 vesicles in the *Klp64D*^*K5*^ homozygous background (Dey et al., 2017), the anterograde movement of Rab4 vesicles was moderately affected in the cholinergic neurons in the heterozygous (*Klp64D*^*K5*^*/+*) background (Fig. 5C-E). We noted that there was a significant increase in the pool of stationary vesicles (Fig. 5C; n≥14 segmental nerves, N≥3 larvae), suggesting a major role of kinesin-2 in initiation of Rab4 vesicle transport.

As expected (Dey et al., 2017), this was also accompanied by a significant reduction of the anterograde segmental velocity and run length (Fig. 5D-E; n>150 runs, N=3 larvae), suggesting that *Klp64D*^*K5*^ mutation could act as a dominant negative allele and suppress the kinesin-2 function. However, acute insulin stimulation in the *Klp64D*^*K5*^*/+* background rescued the motility in the stationary pool of vesicles, enhanced the anterograde flux to the expected level (Fig. 5C; n≥21 segmental nerves, N≥3 larvae) and significantly increase the frequency of fast-moving anterograde runs (Fig. 5D and Table S1; n>150 runs, N=3 larvae).

Altogether, these results suggested that while kinesin-2 plays a critical role in promoting the Rab4 vesicle transport, a different anterograde motor is likely involved in activating anterograde transport and increasing the anterograde flux of Rab4 vesicles downstream of insulin signaling in the axons (Fig. 5F). Therefore, unlike the adipocytes, either insulin stimulation does not promote kinesin-2 recruitment on Rab4 vesicles in the axons or if recruited, does not result in any alterations to the net flux of the Rab4 vesicle movement.

### Klp98A recruitment via PI(3)P-mediated lipid signaling activates anterograde transport of Rab4 vesicles downstream of insulin stimulation

Using a candidate-based screen in rat embryonic fibroblasts, we have previously shown that kinesin-2 (KIF3A) and kinesin-3 (KIF13A/B) likely interact with Rab4 vesicles (Dey et al., 2017). Further validation using genetic perturbations had also revealed that kinesin-2 is the major motor responsible for anterograde transport of Rab4 vesicles (Dey et al., 2017). However, a lack of evidence for kinesin-2 acting as an effector downstream to insulin signaling in activating the anterograde transport of Rab4 vesicles in the axons (Fig. 5C) prompted us to reinvestigate the subject.

With all the previous results, we reasoned that PI(3)P-mediated lipid signaling was one of the strongest hooks that was important for activating kinesin-mediated anterograde transport of Rab4 vesicles. Therefore, we looked for a kinesin with a PX/FYVE domain involved in PI(3)P-binding. KIF16B, a kinesin-3 family motor, has been shown to regulate anterograde movement of Rab5 vesicles involved in receptor recycling through early endosomes in HeLa cells(Hoepfner et al., 2005). Though KIF16B was suggested to be absent from axons by a previous study (Farkhondeh et al., 2015), we tested whether Klp98A – *Drosophila* homolog of KIF16B – could accelerate the movement of Rab4 vesicles downstream to insulin signaling. First, we demonstrated that ectopically expressed Klp98AGFP comigrated with Rab4mRFP vesicles in the axons (Fig. 6A and Supplemental Movie S11; N=2). Next, cell-specific Klp98A knockdown significantly reduced the anterograde flux, segmental velocity and run length of Rab4 vesicles (Fig. 6B-D; n≥9 segmental nerves, n>200runs, N≥4 larvae). Unlike what was observed with perturbations of kinesin-2 function (Fig. 5C), acute insulin stimulation did not rescue the acute anterograde flux, velocity, and run length defects of Rab4 vesicles in Klp98A RNAi background (Fig. 6B-D; n≥9 segmental nerves, n>200runs, N≥4 larvae). Altogether, these observations suggest that insulin stimulation could help recruit Klp98A onto the Rab4 vesicles through Vps34 mediated PI(3)P formation. To test this hypothesis, we estimated the number of Rab4mRFP vesicles labelled with the Klp98AGFP in segmental nerve axons. It revealed that insulin stimulation could significantly increase the frequency of Klp98AGFP localization on Rab4 vesicles and this increase was completely abrogated in the presence of Vps34-specific inhibitor (Fig. 6E-F; n≥10 segmental nerves, N≥3 larvae, >1800 total vesicles). Molecularly, our data demonstrates that Klp98A recruitment on Rab4 vesicles is regulated by the insulin-dependent activation of Vps34 (Fig. 6G) and is a critical step in increasing the anterograde velocity, processivity and flux of Rab4 vesicles in the axons of cholinergic neurons of *Drosophila*. Additionally, our results also suggest that kinesin-2 could play an important role in initiating anterograde movement of Rab4 vesicles (Fig. 5C). In effect, a concerted action of these two kinesin motors in the axon could propel the Rab4 vesicles towards the synapse. Lastly, in view of the previous reports (Dey et al., 2017; Imamura et al., 2003), our data also suggests that Rab4 activity would play a crucial role in recruiting both these kinesin motors on Rab4 vesicles.

**Figure 6:**
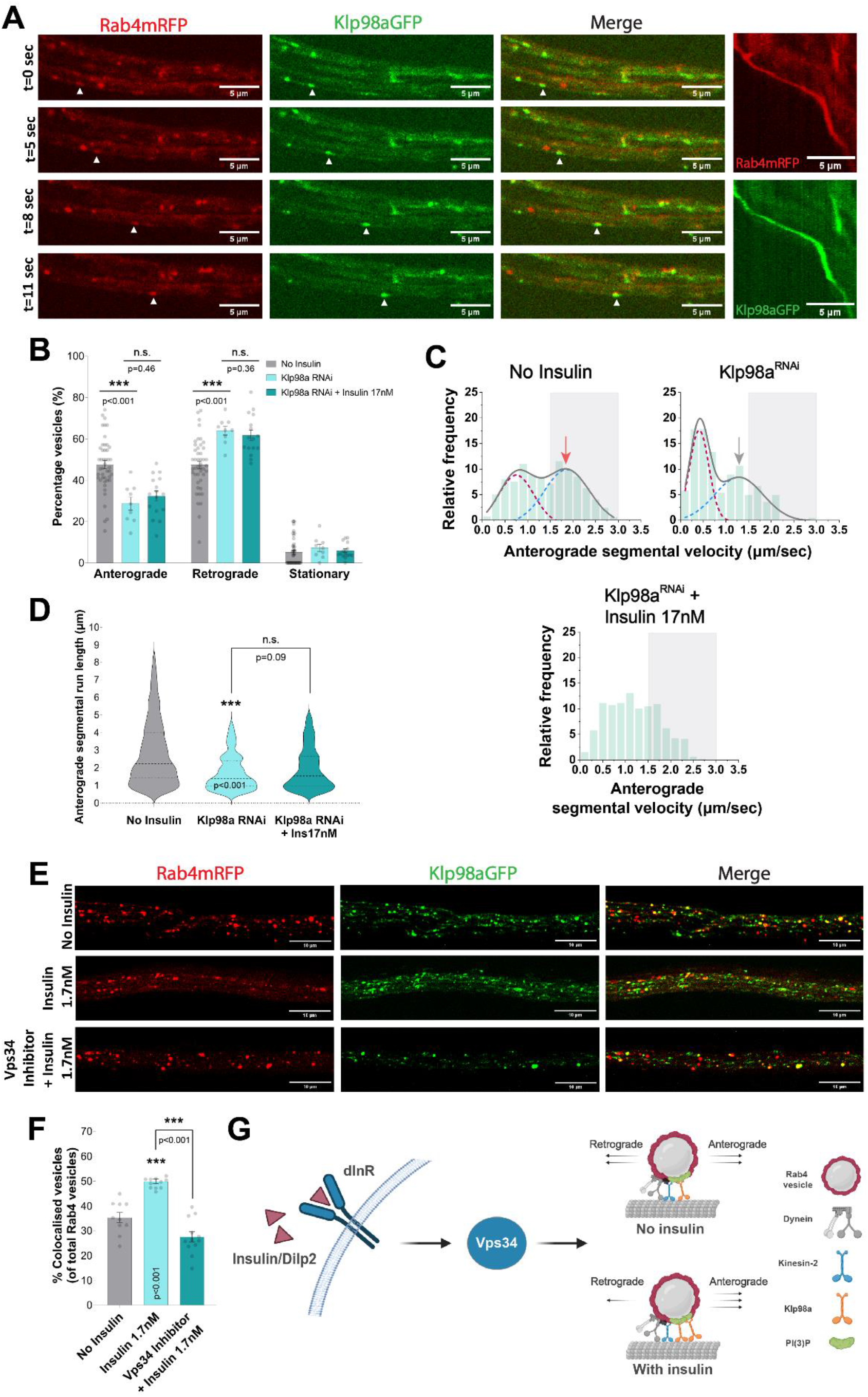
Klp98A recruitment through a Vps34/PI(3)P-dependent pathway regulates the increase in anterograde flux of Rab4 vesicles downstream of insulin signaling. **A)** Time-lapse images of segmental nerve axons depicting comigration of Rab4 vesicles (marked in red) and Klp98A, KIF16B ortholog (marked in green). Kymographs at the right corner illustrate the movement of Rab4 vesicles with Klp98A (see Supplemental Movie S11). **B)** The fraction of Rab4 vesicles moving in the net anterograde, retrograde, and stationary groups based on net total displacement across different genetic and pharmacological perturbations (see Supplemental Movie S6). All statistical comparisons are done using Mann-Whitney U test. **C)** Frequency histograms depicting anterograde segmental velocity (µm/sec) of Rab4 vesicles across different treatment groups. Grey solid lines indicate the cumulative fit of bimodal distribution while the red and blue dashed lines indicate the individual gaussian distributions from the bimodal histogram. Curve fitting has been omitted in one group due to lack of an optimal fit and the fast-moving anterograde runs (1.5 - 3.0 µm/sec) have been marked with a gray box. The red and grey arrows indicate the peak value in the fast-moving range for the control and test group(s), respectively. Comparisons to the ‘No Insulin’ control group are done using Kolmogorov-Smirnov (KS) test. See also Table S1. **D)** Violin plots depicting anterograde segmental run lengths (µm) of Rab4 vesicles across different genetic and pharmacological perturbations. All statistical comparisons are to the ‘No Insulin’ control group (unless specified otherwise) and are done using Mann-Whitney U test. **E)** Representative images from different acute pharmacological perturbations (for 15 minutes) post-fixation showing Rab4-mRFP vesicles (marked in red) and Klp98A-GFP vesicles (marked in green) in segmental nerve axons of cholinergic neurons. **F)** Percentage of Rab4-mRFP and Klp98A-GFP colocalized vesicles across different pharmacological perturbations. All statistical comparisons are to the ‘No Insulin’ control group and are done using Mann-Whitney U test. **G)** Schematic depicting a summary of signaling cascade where activation of insulin signaling via Dilp2-InR-Vps34 axis recruits the PI(3)P-binding kinesin-3 motor Klp98A on Rab4 vesicles and increases their anterograde flux.

## Discussion

### Effects of neuronal insulin signaling on Rab4-associated recycling endosomes in axons

Endosomal trafficking and recycling in neurons are essential for neuronal development and survival (Schmidt & Haucke, 2007; Yap & Winckler, 2012). Directed flux of endosomal traffic is a critical regulator of several neuronal processes like signaling, autophagy, synaptic vesicle recycling, and neurotransmission (Liu et al., 2022; Olenick et al., 2019; Zhao & Zhang, 2019). However, signaling cues regulating directed endosomal flux in the axons remain relatively understudied. In this context, we show that fine-tuning insulin signaling could serve as a cue to regulate the directionality and flux of overall traffic of Rab4 vesicles in the axons.

Insulin signaling in the *Drosophila* brain has been implicated in age related cognitive decline. Besides this, insulin signaling in neurons has been shown to regulate synaptic density (Chiu et al., 2008), neurodegeneration (Schubert et al., 2004), aging (Kenyon, 2010), and behavior (Kleinridders & Pothos, 2019) in various other model systems. However, its effect on long-range axonal transport, which is essential for maintenance of neuronal homeostasis, remained unknown. Here, we show that signaling via insulin/Dilp2 and dInR could accelerate the anterograde movement of a subset of Rab4 vesicles in the cholinergic neurons of *Drosophila*. This observation could have significant physiological bearings as increased accumulation of activated Rab4 at the presynaptic terminals is known to adversely affect synaptic density in *Drosophila* CNS (Dey et al., 2017), and elevated Rab4 levels in both CBF and hippocampal CA1 neurons are positively correlated with neurodegeneration and cognitive decline (Ginsberg et al., 2010, 2011). Our results, together with other findings suggest that insulin signaling could manifest its effect on synapse organization, neurodegeneration, and aging, in-part via altered trafficking of Rab4 endosomes.

### Insulin signaling preferentially activates Vps34 and increases PI3P on Rab4 vesicles in the axons

Insulin signaling is known to activate PI3Kinases, particularly the Class I PI3K in peripheral tissues (Boucher et al., 2014), which activates Rab4 and early endosomal recycling. Here, we show that insulin stimulation preferentially activates Class-III PI3K, also known as Vps34, which is involved in accelerating the speed of Rab4 vesicles. Thus, the mechanism activating the Rab4 vesicle trafficking downstream of the insulin receptor appears to be quite distinct in the axons as compared to peripheral tissues. Vps34 – the lipid kinase that exclusively produces PI(3)P inside the cell – plays a key role in regulating neuronal processes like autophagy (Juhász et al., 2008), synaptic vesicle recycling (Cremona et al., 1999), neurotransmission (K. Li et al., 2019), and neuronal survival (McKnight et al., 2014). Recent studies have also shown that PI(3)P-positive endosomes have a critical role in regulating synaptic vesicle recycling and neurotransmission (Liu et al., 2022). In addition, recent studies have also shown that nutrient-sensitive PI(3)P-mediated lipid signaling on endosomes regulates ER shape and mitochondrial function (Jang et al., 2022). However, the source, identity, and mechanism(s) regulating PI(3)P formation on endosomes in these studies remains unclear. Our study shows that neuronal insulin signaling promotes Vps34-dependent activation of anterograde transport of Rab4-PI(3)P vesicles, which could also act as an active source of replenishing the pool of PI(3)P vesicles enriched at the synapses.

### Klp98A recruitment on Rab4 vesicles and its implications on selective augmentation of Rab4 vesicle flux towards the synapse

Klp98A, the *Drosophila* ortholog of mammalian KIF16B, is a PI(3)P-binding, kinesin-3 family motor protein. KIF16B is recruited on the vesicles through a direct interaction with Rab14 (Ueno et al., 2011). Here, for the first time, we show the recruitment of Klp98A motor onto Rab4 vesicles in the axons which increases significantly due to acute insulin stimulation. KIF16B has previously been shown to bind to PI(3)P through PX domain in its C-terminal tail region (Hoepfner et al., 2005). In this context, the results described here provide an insight into this process by linking Vps34 activity to Klp98A/KIF16B recruitment on Rab4 vesicles. We also show that Vps34 activity regulates the level of PI(3)P and Klp98A motor on Rab4 vesicles, which finally regulates their velocity and anterograde flux. Lastly, an unbiased proteomic screen to identify all Rab4 effectors identified Vps34 as one of the Rab4 effectors (Gillingham et al., 2014), suggesting a potential recruitment mechanism for Vps34 on Rab4 vesicles for the local production of PI(3)P. Together, this indicates that the Vps34 activation downstream of insulin signaling in the axon could critically determine the extent of Rab4 vesicle flux towards the synapse.

A previous study in mouse hippocampal neurons showed that KIF16B is required for somatodendritic localization of early endosomes which helps in trafficking of AMPA and NGF receptors (Farkhondeh et al., 2015). However, whether there is any role of KIF16B in regulating long-range axonal transport *in vivo* remained unclear. We found that Klp98A regulates anterograde axonal transport of Rab4-associated endosomes in *Drosophila* cholinergic neurons. In addition, we also noted that not all motile Klp98A-positive vesicles in axons have Rab4, indicating that Klp98A transports other cargos apart from Rab4 vesicles in axons. Interestingly, mutations in the cargo-binding PX domain of KIF16B have recently been reported in patients with intellectual disability syndrome (Alsahli et al., 2018), although the underlying cause remains to be investigated. Thus, our results could provide a critical mechanistic input because perturbations in Rab4 levels and functionality also affect synaptic homeostasis, which happens to be one of the hallmarks of intellectual disabilities (Zoghbi & Bear, 2012). Therefore, it will also be worth investigating in the future if there is a synergistic interaction between Rab4 and Klp98A keeping in mind their overlapping functions and phenotypes in the CNS.

Cell biological studies suggested that KIF16B drives the fission of early endosome to form tubules (Skjeldal et al., 2012), receptor recycling and degradation(Hoepfner et al., 2005), transcytosis (Perez Bay et al., 2013), and polarized transport of growth factor receptors (Ueno et al., 2011). In addition, the Klp98A motor is implicated in the maturation of autophagic vesicles by promoting fusion through a motor-independent function (Mauvezin et al., 2016). Further, formation of PI(3)P on Rab11 vesicles upon starvation has been shown to serve as a platform for autophagosome formation in HeLa cells (Puri et al., 2018). However, such platforms and/or the identity of PI(3)P membranes required for autophagosome formation in neurons remains unknown. In this context, it may be conjectured that the activation of Vps34-PI3P-Klp98A-dependent movement of Rab4 vesicles towards the synapse in *Drosophila*, as revealed by the above data, could serve as a platform for regulating autophagosome formation at the synapse and reduce the overall synaptic volume. This supposition is also consistent with the results indicating that constitutive activation of Rab4 in cholinergic neurons could increase the Rab4 enrichment at the synapse and reduce the density of synaptic junctions (Dey et al., 2017).

## METHODS

### Larval fillet preparation: Time-lapse imaging and fixed tissue preparations

Larvae were dissected in 1x HL3.1 buffer (Feng et al., 2004) (containing 70 mM NaCl, 5 mM KCl, 1.5 mM CaCl_2_, 4 mM MgCl_2_, 10 mM NaHCO_3_, 5 mM Trehalose, 115 mM Sucrose, and 5 mM HEPES) and insulin (Sigma-Aldrich), Dilps (Pheonix Pharmaceuticals Inc.), PI3K inhibitors (Sigma-Aldrich) were finally reconstituted in HL3.1 buffer for treatment followed by mounting on the coverslip cavity chamber. A small piece of tissue drenched in water was placed on the side of the petri-dish to maintain humidity. The dish is then covered and sealed with Parafilm and immediately taken for time-lapse imaging. Epifluorescence and spinning disc confocal (SDC) time-lapse imaging was performed using Nikon Ti Eclipse TIRF Microscope (installed with a Yokogawa CSU-W1 disk with pinhole size 50 µm for SDC microscopy) operated with running Nikon Elements software using a 100x (1.4 NA) oil-immersion objective (binning=1) with Andor iXon EMCCD camera and sCMOS camera (Zyla 4.2 plus sCMOS) respectively.

Live movies were recorded for all conditions at a frame rate of 8-10 frames per second (fps). For fix tissue analysis, fillet preparations were immediately fixed in 4% paraformaldehyde in 1X PBS for 15-20 minutes at room temperature (RT). Following fixation, they were rinsed thrice with 1X PBS and mounted on a coverslip with a drop of VECTASHIELD® antifade mounting medium (Vector laboratories).

### Insulin/Dilp and drug treatments

Stock solutions of Insulin (Sigma, I0516) and Dilp2/5 (Pheonix Pharmaceuticals Inc., 036-17 & 035-96) were diluted and prepared in HL3.1 buffer, respectively. Likewise, stock solutions of non-specific PI3K inhibitor (5mM) LY294002 (Abcam, #ab120243), HS173 (50µM) (Sigma Aldrich, 5.32384), SAR405 (5mM) (Sigma-Aldrich, 5.33063) were prepared in DMSO (Sigma) and used at effective concentrations of 50µM, 50nM, and 25µM reconstituted in HL3.1 buffer respectively.

### Quantification of axonal transport and statistical analyses

All live-imaging movies were analysed on ImageJ/Fiji® (https://imagej.net/Fiji). KymoAnalyzer plugin(Neumann et al., 2017) was used for estimation of axonal transport parameters like velocity, run length, and cargo population. Input pixel size and frame rate were used as per the movies recorded.

### Statistical information

Origin™ 2019 version 9.60 was used for plotting the frequency histograms (bin size 0.2 µm/sec) and curve fitting was done using Multiple Peak Fit with Gaussian Peak Function. GraphPad Prism 9.5.1 for windows was used for plotting scatter plots and calculation of the Pearson coefficient (r). Nonparametric Kolmogorov-Smirnov test for comparing population distributions and Mann-Whitney U tests were performed using Origin™ 2020 and GraphPad Prism, respectively. The statistical tests, ‘N’, and ‘n’ values are specified in the main text/figure legends for all figures.

### Colocalization estimation

Percentage of colocalized vesicles (Fig 5, 6) were estimated using a standard automated ImageJ plugin ComDet v0.5.5 developed by Eugene Katrukha from Utrecht University https://github.com/UU-cellbiology/ComDet). Imaging parameters for ComDet were chosen such that we could detect >95% vesicles in each image (adjusting pixel size, intensity threshold, and distance between colocalised spots) and automated analysis was validated using manual counting for the control dataset.

## ACKNOWLEDGEMENTS

We thank all K.R. and Howard lab members for all the help and technical support. We also thank Sundar Ram Naganathan and Varsha Mahapatra for comments on the manuscript. We acknowledge the early vital inputs from Komal Raina in the project; Emmanuel Derivery for sharing Klp98A fly stocks and discussions; Swagata Dey and Michael Hanna for discussions; Nishka Malde for assistance in generation of recombinant fly stock; Special thanks to K.V. Boby for maintaining the microscopy facility at DBS, TIFR. J.H. and S.S. were supported by the NIH grant R01 NS118884 and the NSF grant GR117147. The research work at Yale University was also supported by Fulbright-Nehru Doctoral Research Fellowship 2751/FDNR/2022-2023 (to K.S.). The overall research was funded by DST SERB CRG/2020/005396, an intramural grant to K.R., and a fellowship to K.S. and S.D. from TIFR, DAE, Government of India.

## AUTHOR CONTRIBUTIONS

Experimentation: K.S. and S.D.; conceptualization: K.S., S.D., and K.R.; formal analysis and validation: K.S., S.D., S.S., J.H., and K.R.; visualization and figure preparation: K.S., S.D., and K.R.; writing – original draft: K.S. and K.R.; writing – review and editing: K.S., S.D., S.S., J.H., and K.R.; funding acquisition: K.S., K.R., and J.H.; supervision: K.R. and J.H. (at Yale).

## Supplementary Figures

**Figure S1.**
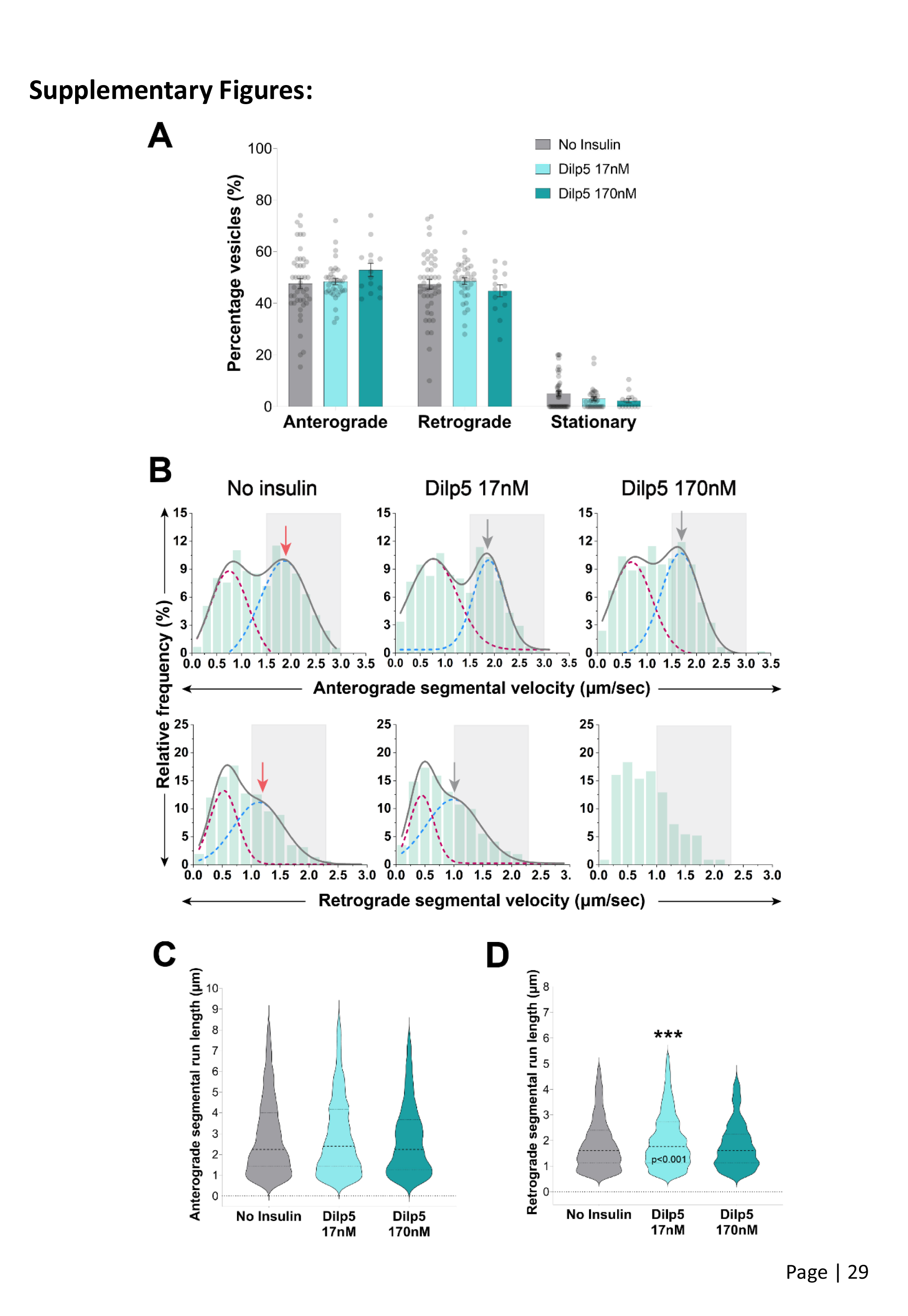
(Linked to Figure 1): Treatment with Dilp5 has no effect on overall flux of Rab4 vesicles in *Drosophila* cholinergic neurons. **A)** Percentage of Rab4 vesicles divided in the net anterograde, retrograde, and stationary groups based on the net displacement (in 1minute) across different treatment groups. All statistical comparisons are to the ‘No Insulin’ control group and are done using Mann-Whitney U test (Supplemental Movie S7). **B)** Frequency histograms of anterograde (top row) and retrograde (bottom row) segmental velocity (µm/sec) of Rab4 vesicles from different treatment groups. Grey solid lines indicate the cumulative fit of bimodal distribution while the red and blue dashed lines indicate the individual gaussian distributions. Curve fitting has been omitted in one group due to lack of an optimal fit and the fast-moving runs are marked with a gray box in both the anterograde (1.5 - 3.0 µm/sec) and retrograde (1.0 - 2.25 µm/sec) velocity plots. The red and grey arrows indicate the peak value of the fast-moving range for the control and test group(s), respectively. Comparisons to the ‘No Insulin’ control group are done using Kolmogorov-Smirnov (KS) test. **C and D)** Violin plots depicting anterograde (C) and retrograde (D) segmental run lengths (µm) of Rab4 vesicles across different treatment groups. All statistical comparisons are to the ‘No Insulin’ control group and are done using Mann-Whitney U test. (p values are mentioned on the plot in cases where the comparison is statistically significant.)

**Figure S2.**
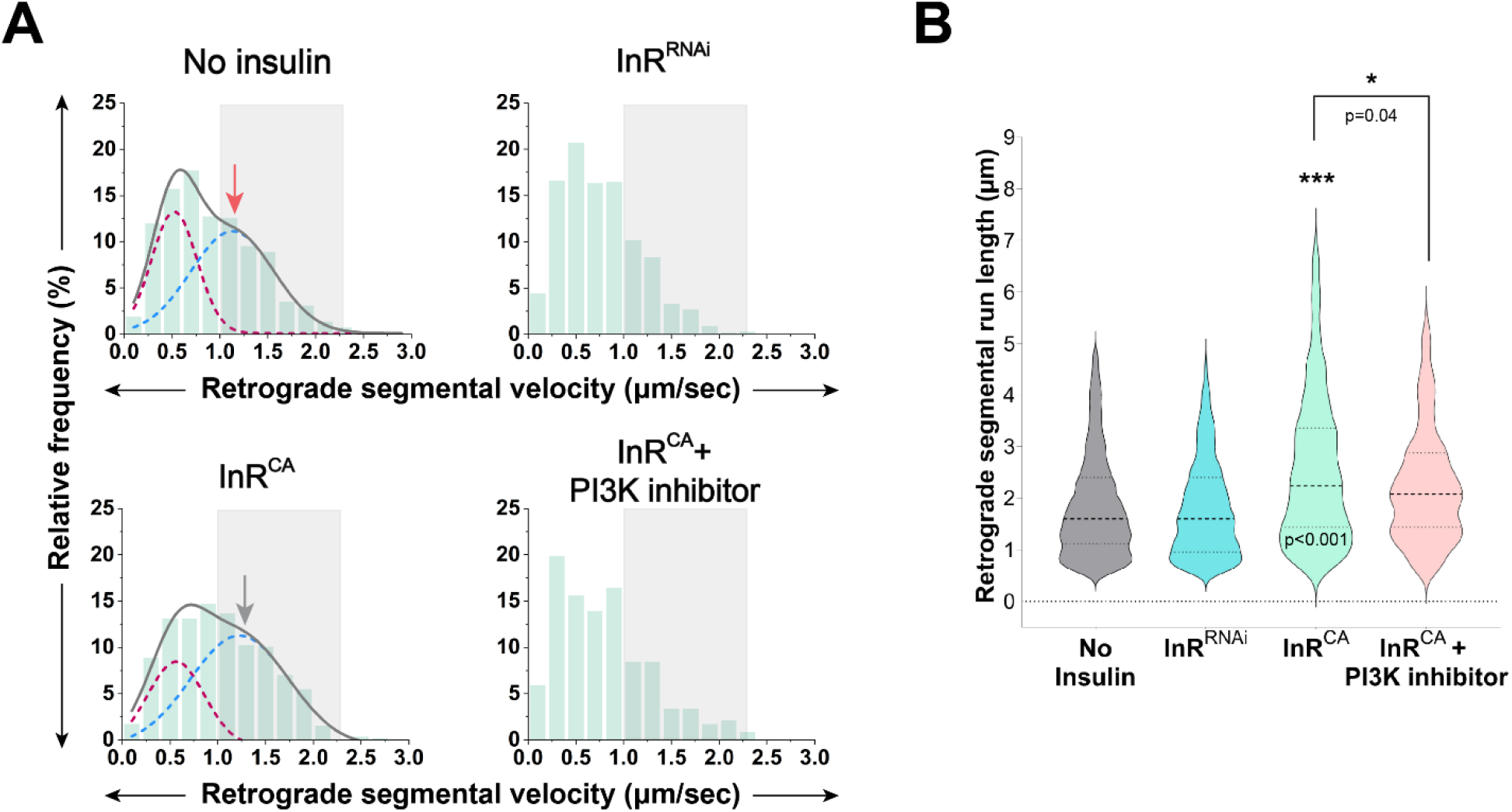
(Linked to Figure 2): Perturbation of insulin signaling also affects retrograde transport parameters of Rab4 vesicles in the axons. **A)** Frequency histograms depicting retrograde segmental velocity (µm/sec) of Rab4 vesicles across different genetic and pharmacological perturbations (see Supplemental Movie S2). Grey solid lines indicate the cumulative fit while the red and blue dashed lines indicate the individual gaussian distributions from the bimodal curve. Curve fitting has been omitted in certain groups due to lack of an optimal fit and the fast-moving retrograde runs (1.0 - 2.25 µm/sec) have been marked with a gray box in all the velocity plot. The red and grey arrows indicate the peak value in the fast-moving range for the control and test group(s), respectively. Comparisons to the ‘No Insulin’ control group are done using Kolmogorov-Smirnov (KS) test. **B)** Violin plots depicting retrograde segmental run lengths (µm) of Rab4 vesicles from different groups. All statistical comparisons are to the ‘No Insulin’ control group unless specified otherwise and are done using Mann-Whitney U test. (p values are mentioned on the plot only in cases where the comparison is statistically significant.)

**Figure S3.**
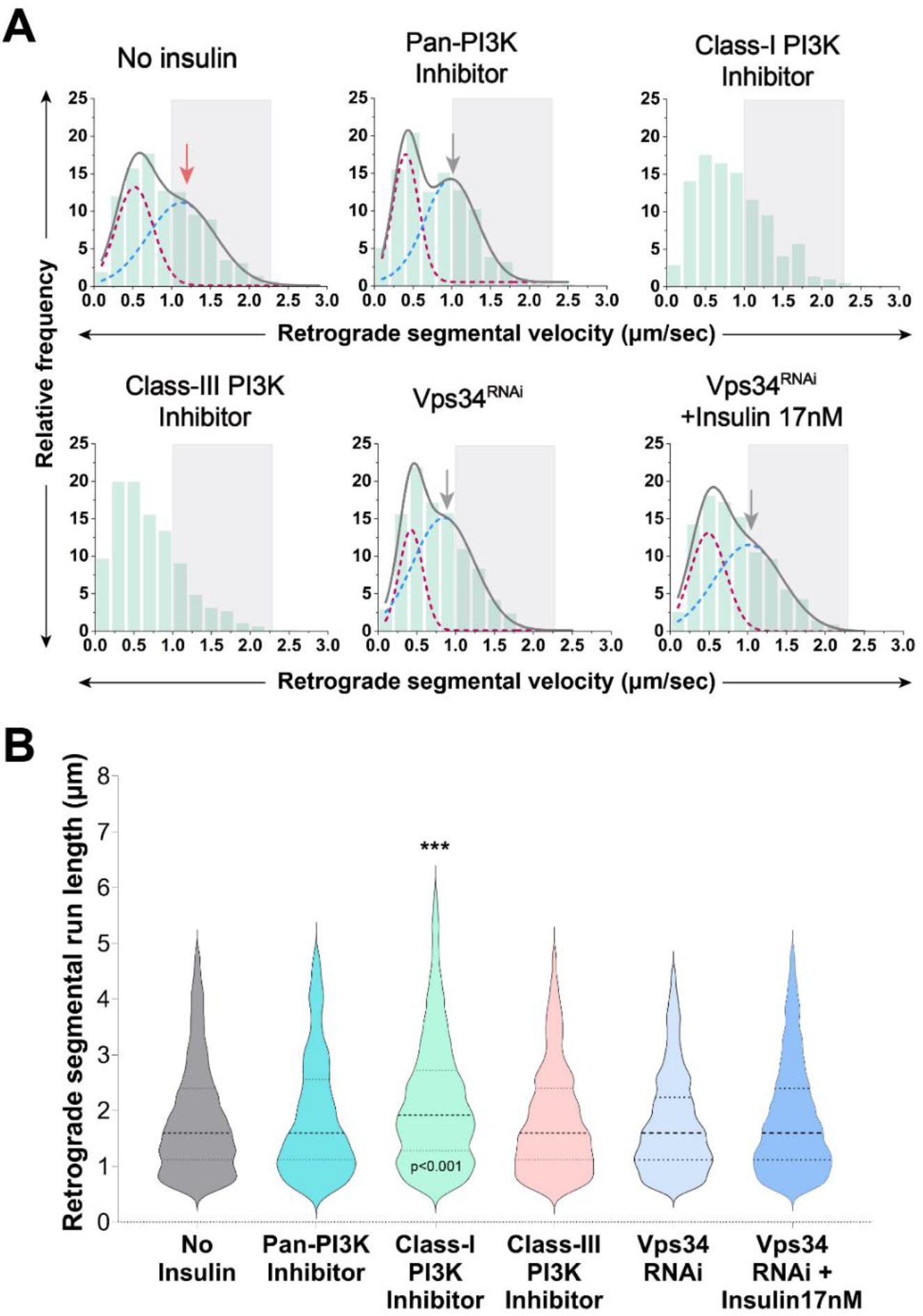
(Linked to Figure 3): Acute inhibition and knockdown of Class-III PI3K/Vps34 reduces the retrograde velocity but has no effect on retrograde run length. **A)** Frequency histograms depicting retrograde segmental velocity (µm/sec) of Rab4 vesicles across different genetic and pharmacological perturbations (see Supplemental Movie S3-4). Grey solid lines indicate the cumulative fit of bimodal distribution while the red and blue dashed lines indicate the individual gaussian distributions from the bimodal curve. Curve fitting has been omitted in certain groups due to lack of an optimal fit and the fast-moving retrograde runs (1.0 - 2.25 µm/sec) have been marked with a gray box in all the velocity plots. The red and grey arrows indicate the peak value in the fast-moving range for the control and test group(s), respectively. Comparisons to the ‘No Insulin’ control group are done using Kolmogorov-Smirnov (KS) test. **B)** Violin plots depicting retrograde segmental run lengths (µm) of Rab4 vesicles across different genetic and pharmacological perturbations. All statistical comparisons are to the ‘No Insulin’ control group (unless specified otherwise) and are done using Mann-Whitney U test. (p values are mentioned on the plot only in cases where the comparison is statistically significant.)

**Figure S4.**
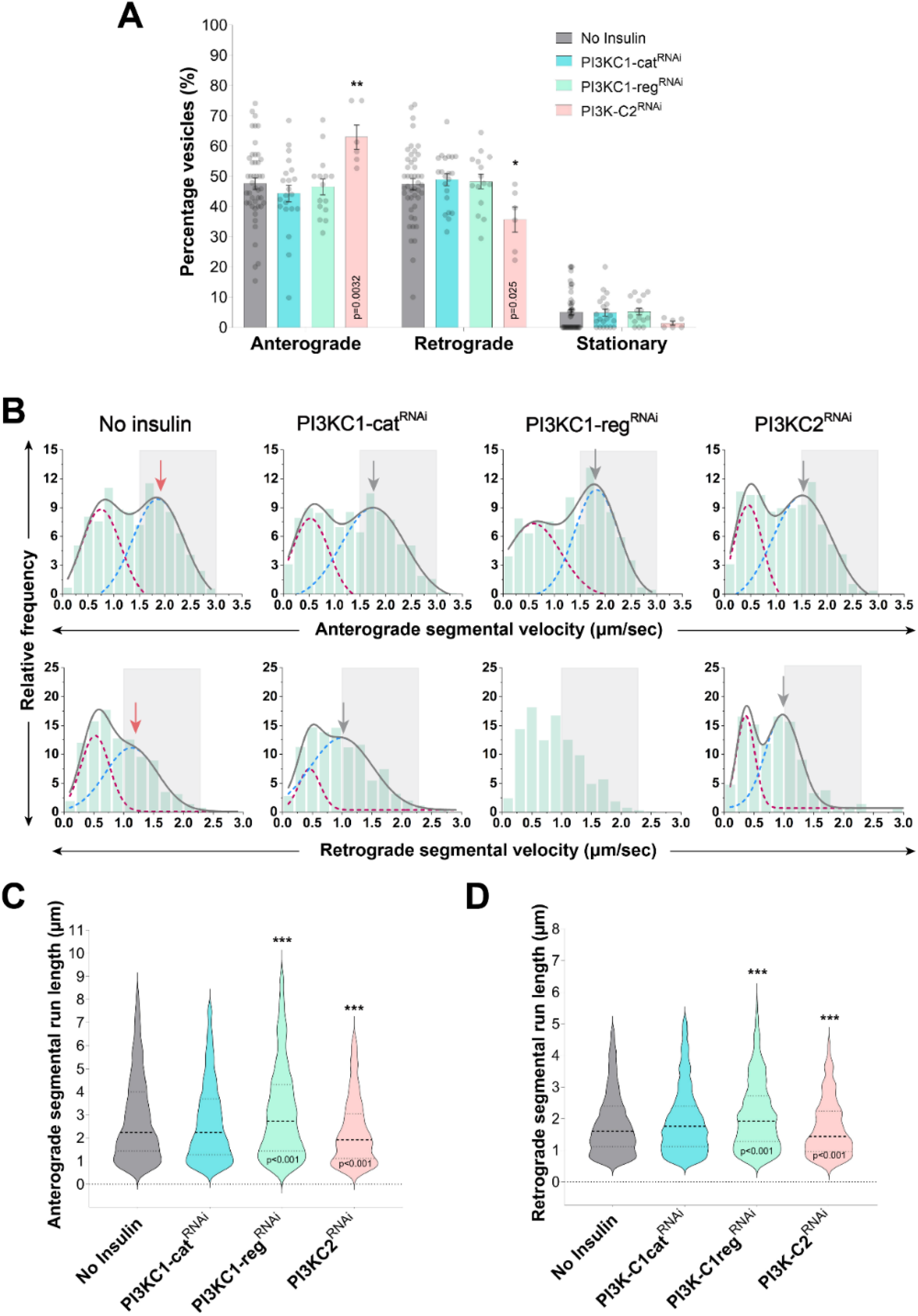
(Linked to Figure 3): Knockdown of Class-I and Class-II PI3Ks do not have a significant effect on anterograde flux of Rab4 vesicles in the axons. **A)** Percentage of Rab4 vesicles divided in the net anterograde, retrograde, and stationary groups based on the net displacement (in 1minute) across different genetic perturbations (see Supplemental Movie S8). All statistical comparisons are to the ‘No Insulin’ control group and are done using Mann-Whitney U test. **B)** Frequency histograms depicting anterograde (top row) and retrograde (bottom row) segmental velocity (µm/sec) of Rab4 vesicles across different genetic perturbations. Grey solid lines indicate the cumulative fit of bimodal distribution while the maroon and blue dashed lines indicate individual gaussian distributions. The fast-moving runs have been marked with a gray box in both anterograde (1.5 - 3.0 µm/sec) and retrograde (1.0 - 2.25 µm/sec) velocity histograms. The red and grey arrows indicate the peak value in the fast-moving range for the control and test group(s), respectively. Comparisons to the ‘No Insulin’ control group are done using Kolmogorov-Smirnov (KS) test. See also Table S1. **C and D)** Violin plots depicting anterograde (C) and retrograde (D) segmental run lengths (µm) of Rab4 vesicles across different genetic perturbations. All statistical comparisons are to the ‘No Insulin’ control group and are done using Mann-Whitney U test. (p values are mentioned on the plot only in cases where the comparison is statistically significant.)

**Figure S5.**
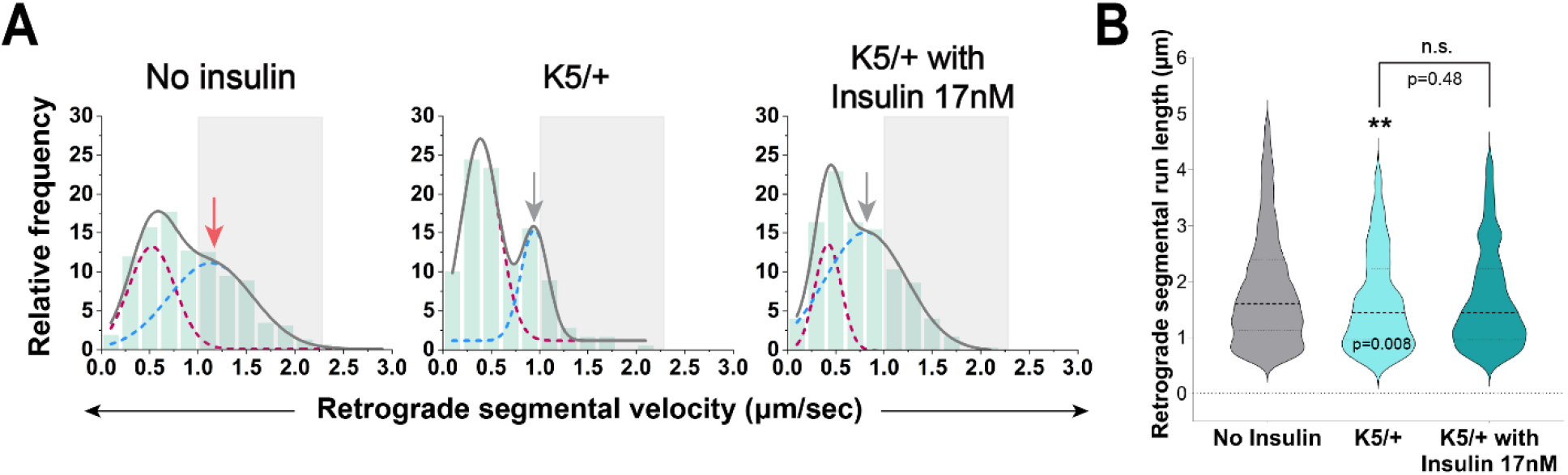
(Linked to Figure 5): Heterozygous mutant of Klp64D^K5^ affects retrograde transport parameters of Rab4 vesicles in cholinergic neurons. **A)** Frequency histograms depicting retrograde segmental velocity (µm/sec) of Rab4 vesicles across different genetic and pharmacological perturbations. Grey solid lines indicate the cumulative fit of bimodal distribution while the red and blue dashed lines indicate the individual gaussian distributions from the bimodal curve. The fast-moving retrograde runs (1.0 - 2.25 µm/sec) is marked with a gray box in all the velocity plots and the red and grey arrows indicate the peak value in the fast-moving range for the control and test group(s), respectively. Comparisons to the ‘No Insulin’ control group are done using Kolmogorov-Smirnov (KS) test. See also Table S1. **B)** Violin plots depicting retrograde segmental run lengths (µm) of Rab4 vesicles across different genetic and pharmacological perturbations. All statistical comparisons are to the ‘no insulin’ control group (unless specified otherwise) and are done using Mann-Whitney U test. (p values are mentioned on the plot only in cases where the comparison is statistically significant.)

**Figure S6.**
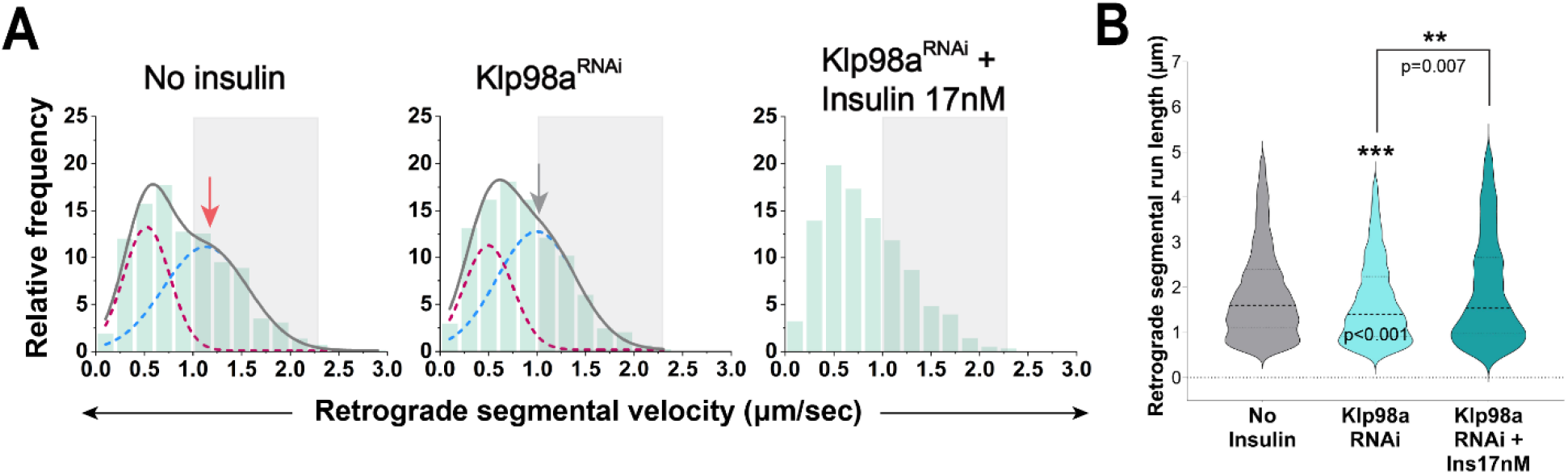
(Linked to Figure 6): Klp98A knockdown also affects retrograde transport parameters of Rab4 vesicles in cholinergic neurons. **A)** Frequency histograms depicting retrograde segmental velocity (µm/sec) of Rab4 vesicles across different genetic and pharmacological perturbations. Grey solid lines indicate the cumulative fit of bimodal distribution while the red and blue dashed lines indicate the individual gaussian distributions from the bimodal curve. The fast-moving retrograde runs (1.0 - 2.25 µm/sec) have been marked with a gray box in all the velocity plots and the red and grey arrows indicate the peak value in the fast-moving range for the control and test group(s), respectively. Comparisons to the ‘No Insulin’ control group are done using Kolmogorov-Smirnov (KS) test (see also Table S1). **B)** Violin plots depicting retrograde segmental run lengths (µm) of Rab4 vesicles across different genetic and pharmacological perturbations. All statistical comparisons are to the ‘No Insulin’ control group (unless specified otherwise) and are done using Mann-Whitney U test. (p values are mentioned on the plot only in cases where the comparison is statistically significant)

**Table S1:** Percentage of fast- and slow-moving vesicles in anterograde and retrograde directions across all pharmacological treatments and genetic perturbations.

| Treatment/Perturbation | Anterograde (%) |  | Retrograde (%) |  |
| --- | --- | --- | --- | --- |
|  | 0-1.6µm/sec | >1.6µm/sec | 0-1µm/sec | >1µm/sec |
| No insulin | 56.82 | 43.18 | 60.05 | 39.95 |
| Insulin 1.7nM | 48.33 | 51.67 | 50.53 | 49.47 |
| Insulin 17nM | 64.1 | 35.9 | 63.05 | 36.95 |
| Dilp2 17nM | 51.34 | 48.66 | 59.98 | 40.02 |
| Dilp2 170nM | 41.46 | 58.54 | 47.83 | 52.16 |
| Dilp5 17nM | 62.32 | 37.68 | 64.63 | 35.37 |
| Dilp5 170nM | 68.83 | 31.17 | 67.42 | 32.58 |
| LY294002 50uM | 78.61 | 21.39 | 68.57 | 31.43 |
| HS173 50nM | 64.07 | 35.93 | 66.15 | 33.85 |
| SAR405 50uM | 81.44 | 18.56 | 78.38 | 21.62 |
| InR<sup>CA</sup> | 42.65 | 57.35 | 51.54 | 48.46 |
| InR<sup>CA</sup> + LY294002 560uM | 70.2 | 29.8 | 71.73 | 28.27 |
| InR<sup>RNAi</sup> | 72.94 | 27.06 | 74.4 | 25.6 |
| Class-I PI3K cat<sup>RNAi</sup> | 59.16 | 40.84 | 57.08 | 42.92 |
| Class-I PI3K reg<sup>RNAi</sup> | 57.13 | 42.87 | 64.04 | 35.96 |
| Class-II PI3K<sup>RNAi</sup> | 70.13 | 29.89 | 64.93 | 35.07 |
| Vps34<sup>RNAi</sup> | 67.52 | 32.48 | 73.31 | 26.69 |
| Vps34<sup>RNAi</sup> + Insulin 17nM | 63.66 | 36.34 | 67.29 | 32.71 |
| Klp64D<sup>K5</sup> | 96.38 | 3.62 | 84.44 | 15.56 |
| Klp64D<sup>K5</sup> + Insulin 17nM | 77.8 | 22.2 | 75.07 | 24.93 |
| Klp98A<sup>RNAi</sup> | 84.44 | 15.56 | 66.41 | 33.59 |
| Klp98A<sup>RNAi</sup> + Insulin 17nM | 73.72 | 26.28 | 68.55 | 31.45 |

## Legends for Movies

**Supplemental Movie S1 (Linked to Figure 1):** Effect of human insulin and Dilp2 treatment on the transport of Rab4mRFP vesicles in the distal axons of *Drosophila* cholinergic neurons. Widefield time-lapse imaging was performed at 10fps and the movies are played at 50fps. Duration of all the representative movies is 30sec.

**Supplemental Movie S2 (Linked to Figure 2):** Time imaging of Rab4 vesicles in the distal axons of *Drosophila* cholinergic neurons across different genetic and pharmacological perturbations. Widefield time-lapse imaging was performed at 10fps and the representative movies are played at 50fps. Duration of all the representative movies is 30sec.

**Supplemental Movie S3 (Linked to Figure 3):** Effect of inhibition of all the PI3Ks on the transport of Rab4mRFP vesicles in the distal axons of *Drosophila* cholinergic neurons using specific chemical inhibitors. Widefield time-lapse imaging was performed at 10fps and the movies are played at 50fps. Duration of all the representative movies is 30sec.

**Supplemental Movie S4 (Linked to Figure 3):** Effect of cell-specific knockdown of Vps34 on the transport of Rab4mRFP vesicles in the distal axons of *Drosophila* cholinergic neurons. Widefield time-lapse imaging was performed at 10fps and the representative movies are played at 50fps. Duration of all the representative movies is 30sec.

**Supplemental Movie S5 (Linked to Figure 5):** Effect of perturbation of kinesin-2 functionality using Klp64D^K5^ allele in a heterozygous background on the transport of Rab4mRFP vesicles in the distal axons of *Drosophila* cholinergic neurons. Widefield time-lapse imaging was performed at 10fps and the representative movies are played at 50fps. Duration of all the representative movies is 30sec.

**Supplemental Movie S6 (Linked to Figure 6):** Effect of cell-specific knockdown of Klp98A on the transport of Rab4mRFP vesicles in the distal axons of *Drosophila* cholinergic neurons. Widefield time-lapse imaging was performed at 8-10fps and the representative movies are played at 50fps. Duration of all the representative movies is 30sec.

**Supplemental Movie S7 (Linked to Figure S1):** Effect of Dilp5 treatment on the transport of Rab4mRFP vesicles in the distal axons of *Drosophila* cholinergic neurons. Widefield time-lapse imaging was performed at 10fps and the movies are played at 50fps. Duration of all the representative movies is 30sec.

**Supplemental Movie S8 (Linked to Figure S4):** Effect of cell-specific knockdown of Class-I and Class-II PI3K on the transport of Rab4mRFP vesicles in the distal axons of *Drosophila* cholinergic neurons. Widefield time-lapse imaging was performed at 8-10fps and the representative movies are played at 50fps. Duration of all the representative movies is 30sec.

**Supplemental Movie S9 (Linked to Figure 5):** Simultaneous dual-color time-lapse imaging performed using Spinning-disc confocal microscopy to visualize comigration of Klp64D-GFP and Rab4mRFP in the distal axons of *Drosophila* cholinergic neurons. Data was collected at 8-9fps and movies are played at 50fps. Duration of the representative movie is 30sec.

**Supplemental Movie S10 (Linked to Figure 4):** Simultaneous dual-color time-lapse imaging performed using Spinning-disc confocal microscopy to visualize comigration of genetically-encoded PI(3)P biosensor (2xFYVE-GFP) and Rab4mRFP in the distal axons of *Drosophila* cholinergic neurons. Data was collected at 8-9fps and movies are played at 50fps. Duration of the representative movie is 20sec.

**Supplemental Movie S11 (Linked to Figure 6:** Simultaneous dual-color time-lapse imaging performed using Spinning-disc confocal microscopy to visualize comigration of Klp98A-GFP and Rab4mRFP in the distal axons of *Drosophila* cholinergic neurons. Data was collected at 8-9fps and movies are played at 50fps. Duration of the representative movie is ∼20sec.

**Supplemental Movie S12 (Linked to Figure 4):** Simultaneous dual-color time-lapse imaging performed using Spinning-disc confocal microscopy to visualize comigration of 2xFYVE-GFP and Rab4mRFP in the distal axons of *Drosophila* cholinergic neurons. Particle 1 (green) and Particle 2 (cyan) are representative examples of particles with a low and high Rab4mRFP/2xFYVE-GFP intensity ratio (as plotted in Figure 4F), respectively. Data was collected at 8-9fps and movies are played at 50fps. Duration of the representative movie is 20sec.

## References

Ahmad, T., Vullhorst, D., Chaudhuri, R., Guardia, C. M., Chaudhary, N., Karavanova, I., Bonifacino, J. S., & Buonanno, A. (2022). Transcytosis and trans-synaptic retention by postsynaptic ErbB4 underlie axonal accumulation of NRG3. Journal of Cell Biology, 221(7), e202110167. 10.1083/jcb.202110167

Akintola, A. A., & Van Heemst, D. (2015). Insulin, Aging, and the Brain: Mechanisms and Implications. Frontiers in Endocrinology, 6. 10.3389/fendo.2015.00013

Alsahli, S., Arold, S. T., Alfares, A., Alhaddad, B., Al Balwi, M., Kamsteeg, E., Al-Twaijri, W., & Alfadhel, M. (2018). KIF16B is a candidate gene for a novel autosomal-recessive intellectual disability syndrome. American Journal of Medical Genetics Part A, 176(7), 1602–1609. 10.1002/ajmg.a.38723

Augustin, H., McGourty, K., Allen, M. J., Madem, S. K., Adcott, J., Kerr, F., Wong, C. T., Vincent, A., Godenschwege, T., Boucrot, E., & Partridge, L. (2017). Reduced insulin signaling maintains electrical transmission in a neural circuit in aging flies. PLOS Biology, 15(9), e2001655. 10.1371/journal.pbio.2001655

Bago, R., Malik, N., Munson, M. J., Prescott, A. R., Davies, P., Sommer, E., Shpiro, N., Ward, R., Cross, D., Ganley, I. G., & Alessi, D. R. (2014). Characterization of VPS34-IN1, a selective inhibitor of Vps34, reveals that the phosphatidylinositol 3-phosphate-binding SGK3 protein kinase is a downstream target of class III phosphoinositide 3-kinase. Biochemical Journal, 463(3), 413–427. 10.1042/BJ20140889

Behesti, H., Fore, T. R., Wu, P., Horn, Z., Leppert, M., Hull, C., & Hatten, M. E. (2018). ASTN2 modulates synaptic strength by trafficking and degradation of surface proteins. Proceedings of the National Academy of Sciences, 115(41). 10.1073/pnas.1809382115

Bielli, A., Thörnqvist, P.-O., Hendrick, A. G., Finn, R., Fitzgerald, K., & McCaffrey, M. W. (2001). The Small GTPase Rab4A Interacts with the Central Region of Cytoplasmic Dynein Light Intermediate Chain-1. Biochemical and Biophysical Research Communications, 281(5), 1141–1153. 10.1006/bbrc.2001.4468

Boucher, J., Kleinridders, A., & Kahn, C. R. (2014). Insulin Receptor Signaling in Normal and Insulin-Resistant States. Cold Spring Harbor Perspectives in Biology, 6(1), a009191–a009191. 10.1101/cshperspect.a009191

Brogiolo, W., Stocker, H., Ikeya, T., Rintelen, F., Fernandez, R., & Hafen, E. (2001). An evolutionarily conserved function of the Drosophila insulin receptor and insulin-like peptides in growth control. Current Biology, 11(4), 213–221. 10.1016/S0960-9822(01)00068-9

Brown, J. B., Boley, N., Eisman, R., May, G. E., Stoiber, M. H., Duff, M. O., Booth, B. W., Wen, J., Park, S., Suzuki, A. M., Wan, K. H., Yu, C., Zhang, D., Carlson, J. W., Cherbas, L., Eads, B. D., Miller, D., Mockaitis, K., Roberts, J., … Celniker, S. E. (2014). Diversity and dynamics of the Drosophila transcriptome. Nature, 512(7515), 393–399. 10.1038/nature12962

Chiu, S.-L., Chen, C.-M., & Cline, H. T. (2008). Insulin receptor signaling regulates synapse number, dendritic plasticity, and circuit function in vivo. Neuron, 58(5), 708–719. 10.1016/j.neuron.2008.04.014

Cremona, O., Di Paolo, G., Wenk, M. R., Lüthi, A., Kim, W. T., Takei, K., Daniell, L., Nemoto, Y., Shears, S. B., Flavell, R. A., McCormick, D. A., & De Camilli, P. (1999). Essential Role of Phosphoinositide Metabolism in Synaptic Vesicle Recycling. Cell, 99(2), 179–188. 10.1016/S0092-8674(00)81649-9

Dey, S., Banker, G., & Ray, K. (2017). Anterograde Transport of Rab4-Associated Vesicles Regulates Synapse Organization in Drosophila. Cell Reports, 18(10), 2452–2463. 10.1016/j.celrep.2017.02.034

Di Paolo, G., & De Camilli, P. (2006). Phosphoinositides in cell regulation and membrane dynamics. Nature, 443(7112), 651–657. 10.1038/nature05185

Dupont, J., & Holzenberger, M. (2003). Biology of insulin-like growth factors in development. Birth Defects Research Part C: Embryo Today: Reviews, 69(4), 257–271. 10.1002/bdrc.10022

Falk, J., Konopacki, F. A., Zivraj, K. H., & Holt, C. E. (2014). Rab5 and Rab4 Regulate Axon Elongation in the Xenopus Visual System. Journal of Neuroscience, 34(2), 373–391. 10.1523/JNEUROSCI.0876-13.2014

Farkhondeh, A., Niwa, S., Takei, Y., & Hirokawa, N. (2015). Characterizing KIF16B in Neurons Reveals a Novel Intramolecular “Stalk Inhibition” Mechanism That Regulates Its Capacity to Potentiate the Selective Somatodendritic Localization of Early Endosomes. The Journal of Neuroscience, 35(12), 5067–5086. 10.1523/JNEUROSCI.4240-14.2015

Feng, Y., Ueda, A., & Wu, C.-F. (2004). A modified minimal hemolymph-like solution, HL3.1, for physiological recordings at the neuromuscular junctions of normal and mutant Drosophila larvae. Journal of Neurogenetics, 18(2), 377–402. 10.1080/01677060490894522

Gillingham, A. K., Sinka, R., Torres, I. L., Lilley, K. S., & Munro, S. (2014). Toward a Comprehensive Map of the Effectors of Rab GTPases. Developmental Cell, 31(3), 358–373. 10.1016/j.devcel.2014.10.007

Gillooly, D. J. (2000). Localization of phosphatidylinositol 3-phosphate in yeast and mammalian cells. The EMBO Journal, 19(17), 4577–4588. 10.1093/emboj/19.17.4577

Ginsberg, S. D., Alldred, M. J., Counts, S. E., Cataldo, A. M., Neve, R. L., Jiang, Y., Wuu, J., Chao, M. V., Mufson, E. J., Nixon, R. A., & Che, S. (2010). Microarray analysis of hippocampal CA1 neurons implicates early endosomal dysfunction during Alzheimer’s disease progression. Biological Psychiatry, 68(10), 885–893. 10.1016/j.biopsych.2010.05.030

Ginsberg, S. D., Mufson, E. J., Alldred, M. J., Counts, S. E., Wuu, J., Nixon, R. A., & Che, S. (2011). Upregulation of select rab GTPases in cholinergic basal forebrain neurons in mild cognitive impairment and Alzheimer’s disease. Journal of Chemical Neuroanatomy, 42(2), 102–110. 10.1016/j.jchemneu.2011.05.012

Gumy, L. F., Katrukha, E. A., Grigoriev, I., Jaarsma, D., Kapitein, L. C., Akhmanova, A., & Hoogenraad, C. C. (2017). MAP2 Defines a Pre-axonal Filtering Zone to Regulate KIF1-versus KIF5-Dependent Cargo Transport in Sensory Neurons. Neuron, 94(2), 347-362.e7. 10.1016/j.neuron.2017.03.046

Hakuno, F., & Takahashi, S.-I. (2018). 40 YEARS OF IGF1: IGF1 receptor signaling pathways. Journal of Molecular Endocrinology, 61(1), T69–T86. 10.1530/JME-17-0311

Hoepfner, S., Severin, F., Cabezas, A., Habermann, B., Runge, A., Gillooly, D., Stenmark, H., & Zerial, M. (2005). Modulation of Receptor Recycling and Degradation by the Endosomal Kinesin KIF16B. Cell, 121(3), 437–450. 10.1016/j.cell.2005.02.017

Hoogenraad, C. C., Popa, I., Futai, K., Sanchez-Martinez, E., Wulf, P. S., van Vlijmen, T., Dortland, B. R., Oorschot, V., Govers, R., Monti, M., Heck, A. J. R., Sheng, M., Klumperman, J., Rehmann, H., Jaarsma, D., Kapitein, L. C., & van der Sluijs, P. (2010). Neuron Specific Rab4 Effector GRASP-1 Coordinates Membrane Specialization and Maturation of Recycling Endosomes. PLoS Biology, 8(1), e1000283. 10.1371/journal.pbio.1000283

Imamura, T., Huang, J., Usui, I., Satoh, H., Bever, J., & Olefsky, J. M. (2003). Insulin-induced GLUT4 translocation involves protein kinase C-lambda-mediated functional coupling between Rab4 and the motor protein kinesin. Molecular and Cellular Biology, 23(14), 4892–4900. 10.1128/mcb.23.14.4892-4900.2003

Jang, W., Puchkov, D., Samsó, P., Liang, Y., Nadler-Holly, M., Sigrist, S. J., Kintscher, U., Liu, F., Mamchaoui, K., Mouly, V., & Haucke, V. (2022). Endosomal lipid signaling reshapes the endoplasmic reticulum to control mitochondrial function. Science, 378(6625), eabq5209. 10.1126/science.abq5209

Jia, D., Soylemez, M., Calvin, G., Bornmann, R., Bryant, J., Hanna, C., Huang, Y.-C., & Deng, W.-M. (2015). A large-scale in vivo RNAi screen to identify genes involved in Notch-mediated follicle cell differentiation and cell cycle switches. Scientific Reports, 5(1), 12328. 10.1038/srep12328

Juhász, G., Hill, J. H., Yan, Y., Sass, M., Baehrecke, E. H., Backer, J. M., & Neufeld, T. P. (2008). The class III PI(3)K Vps34 promotes autophagy and endocytosis but not TOR signaling in Drosophila. Journal of Cell Biology, 181(4), 655–666. 10.1083/jcb.200712051

Kenyon, C. J. (2010). The genetics of ageing. Nature, 464(7288), 504–512. 10.1038/nature08980

Kleinridders, A., & Pothos, E. N. (2019). Impact of Brain Insulin Signaling on Dopamine Function, Food Intake, Reward, and Emotional Behavior. Current Nutrition Reports, 8(2), 83–91. 10.1007/s13668-019-0276-z

Kumar, M., Ojha, S., Rai, P., Joshi, A., Kamat, S. S., & Mallik, R. (2019). Insulin activates intracellular transport of lipid droplets to release triglycerides from the liver. Journal of Cell Biology, 218(11), 3697–3713. 10.1083/jcb.201903102

Li, H., Janssens, J., De Waegeneer, M., Kolluru, S. S., Davie, K., Gardeux, V., Saelens, W., David, F. P. A., Brbić, M., Spanier, K., Leskovec, J., McLaughlin, C. N., Xie, Q., Jones, R. C., Brueckner, K., Shim, J., Tattikota, S. G., Schnorrer, F., Rust, K., … Zinzen, R. P. (2022). Fly Cell Atlas: A single-nucleus transcriptomic atlas of the adult fruit fly. Science, 375(6584), eabk2432. 10.1126/science.abk2432

Li, K., Chen, H.-S., Li, D., Li, H.-H., Wang, J., Jia, L., Wu, P.-F., Long, L.-H., Hu, Z.-L., Chen, J.-G., & Wang, F. (2019). SAR405, a Highly Specific VPS34 Inhibitor, Disrupts Auditory Fear Memory Consolidation of Mice via Facilitation of Inhibitory Neurotransmission in Basolateral Amygdala. Biological Psychiatry, 85(3), 214–225. 10.1016/j.biopsych.2018.07.026

Liu, G., Kochlamazashvili, G., Puchkov, D., Müller, R., Schultz, C., Mackintosh, A. I., Vollweiter, D., Haucke, V., & Soykan, T. (2022). Endosomal phosphatidylinositol 3-phosphate controls synaptic vesicle cycling and neurotransmission. The EMBO Journal, 41(9). 10.15252/embj.2021109352

Man, H.-Y., Lin, J. W., Ju, W. H., Ahmadian, G., Liu, L., Becker, L. E., Sheng, M., & Wang, Y. T. (2000). Regulation of AMPA Receptor–Mediated Synaptic Transmission by Clathrin-Dependent Receptor Internalization. Neuron, 25(3), 649–662. 10.1016/S0896-6273(00)81067-3

Mauvezin, C., Neisch, A. L., Ayala, C. I., Kim, J., Beltrame, A., Braden, C. R., Gardner, M. K., Hays, T. S., & Neufeld, T. P. (2016). Coordination of autophagosome-lysosome fusion and transport by a Klp98A-Rab14 complex. Journal of Cell Science, jcs.175224. 10.1242/jcs.175224

McKnight, N. C., Zhong, Y., Wold, M. S., Gong, S., Phillips, G. R., Dou, Z., Zhao, Y., Heintz, N., Zong, W.-X., & Yue, Z. (2014). Beclin 1 Is Required for Neuron Viability and Regulates Endosome Pathways via the UVRAG-VPS34 Complex. PLoS Genetics, 10(10), e1004626. 10.1371/journal.pgen.1004626

Moretto, E., Miozzo, F., Longatti, A., Bonnet, C., Coussen, F., Jaudon, F., Cingolani, L. A., & Passafaro, M. (2023). The tetraspanin TSPAN5 regulates AMPAR exocytosis by interacting with the AP4 complex. eLife, 12, e76425. 10.7554/eLife.76425

Nässel, D. R., & Broeck, J. V. (2016). Insulin/IGF signaling in Drosophila and other insects: Factors that regulate production, release and post-release action of the insulin-like peptides. Cellular and Molecular Life Sciences, 73(2), 271–290. 10.1007/s00018-015-2063-3

Neumann, S., Chassefeyre, R., Campbell, G. E., & Encalada, S. E. (2017). KymoAnalyzer: A software tool for the quantitative analysis of intracellular transport in neurons: NEUMANN ET AL. Traffic, 18(1), 71–88. 10.1111/tra.12456

Olenick, M. A., Dominguez, R., & Holzbaur, E. L. F. (2019). Dynein activator Hook1 is required for trafficking of BDNF-signaling endosomes in neurons. Journal of Cell Biology, 218(1), 220–233. 10.1083/jcb.201805016

Pattni, K., Jepson, M., Stenmark, H., & Banting, G. (2001). A PtdIns(3)P-specific probe cycles on and off host cell membranes during Salmonella invasion of mammalian cells. Current Biology, 11(20), 1636–1642. 10.1016/S0960-9822(01)00486-9

Perez Bay, A. E., Schreiner, R., Mazzoni, F., Carvajal-Gonzalez, J. M., Gravotta, D., Perret, E., Lehmann Mantaras, G., Zhu, Y.-S., & Rodriguez-Boulan, E. J. (2013). The kinesin KIF16B mediates apical transcytosis of transferrin receptor in AP-1B-deficient epithelia. The EMBO Journal, 32(15), 2125–2139. 10.1038/emboj.2013.130

Phoenix Pharmaceuticals. (n.d.). https://phoenixpeptide.com/products/view/Peptides/036-17

Puri, C., Vicinanza, M., Ashkenazi, A., Gratian, M. J., Zhang, Q., Bento, C. F., Renna, M., Menzies, F. M., & Rubinsztein, D. C. (2018). The RAB11A-Positive Compartment Is a Primary Platform for Autophagosome Assembly Mediated by WIPI2 Recognition of PI3P-RAB11A. Developmental Cell, 45(1), 114-131.e8. 10.1016/j.devcel.2018.03.008

Ray, K., Perez, S. E., Yang, Z., Xu, J., Ritchings, B. W., Steller, H., & Goldstein, L. S. B. (1999). Kinesin-II Is Required for Axonal Transport of Choline Acetyltransferase in Drosophila. The Journal of Cell Biology, 147(3), 507–518. 10.1083/jcb.147.3.507

Sato, T. K., Overduin, M., & Emr, S. D. (2001). Location, Location, Location: Membrane Targeting Directed by PX Domains. Science, 294(5548), 1881–1885. 10.1126/science.1065763

Schmidt, M. R., & Haucke, V. (2007). Recycling endosomes in neuronal membrane traffic. Biology of the Cell, 99(6), 333–342. 10.1042/BC20070007

Schu, P. V., Takegawa, K., Fry, M. J., Stack, J. H., Waterfield, M. D., & Emr, S. D. (1993). Phosphatidylinositol 3-Kinase Encoded by Yeast VPS 34 Gene Essential for Protein Sorting. Science, 260(5104), 88–91. 10.1126/science.8385367

Schubert, M., Gautam, D., Surjo, D., Ueki, K., Baudler, S., Schubert, D., Kondo, T., Alber, J., Galldiks, N., Küstermann, E., Arndt, S., Jacobs, A. H., Krone, W., Kahn, C. R., & Brüning, J. C. (2004). Role for neuronal insulin resistance in neurodegenerative diseases. Proceedings of the National Academy of Sciences, 101(9), 3100–3105. 10.1073/pnas.0308724101

Semaniuk, U., Piskovatska, V., Strilbytska, O., Strutynska, T., Burdyliuk, N., Vaiserman, A., Bubalo, V., Storey, K. B., & Lushchak, O. (2021). Drosophila insulin-like peptides: From expression to functions – a review. Entomologia Experimentalis et Applicata, 169(2), 195–208. 10.1111/eea.12981

Shibata, H., Omata, W., & Kojima, I. (1997). Insulin Stimulates Guanine Nucleotide Exchange on Rab4 via a Wortmannin-sensitive Signaling Pathway in Rat Adipocytes. Journal of Biological Chemistry, 272(23), 14542–14546. 10.1074/jbc.272.23.14542

Shieh, J. C.-C., Huang, P.-T., & Lin, Y.-F. (2020). Alzheimer’s Disease and Diabetes: Insulin Signaling as the Bridge Linking Two Pathologies. Molecular Neurobiology, 57(4), 1966–1977. 10.1007/s12035-019-01858-5

Shin, G. J., Pero, M. E., Hammond, L. A., Burgos, A., Kumar, A., Galindo, S. E., Lucas, T., Bartolini, F., & Grueber, W. B. (2021). Integrins protect sensory neurons in models of paclitaxel-induced peripheral sensory neuropathy. Proceedings of the National Academy of Sciences, 118(15). 10.1073/pnas.2006050118

Sigma. (n.d.). https://www.sigmaaldrich.com/deepweb/assets/sigmaaldrich/product/documents/108/221/i0259pis.pdf

Skjeldal, F. M., Strunze, S., Bergeland, T., Walseng, E., FGregers, T., & Bakke, O. (2012). The fusion of early endosomes induces molecular motor-driven tubule formation and fission. Journal of Cell Science, jcs.092569. 10.1242/jcs.092569

Tokarz, V. L., MacDonald, P. E., & Klip, A. (2018). The cell biology of systemic insulin function. Journal of Cell Biology, 217(7), 2273–2289. 10.1083/jcb.201802095

Tsukazaki, T., Chiang, T. A., Davison, A. F., Attisano, L., & Wrana, J. L. (1998). SARA, a FYVE Domain Protein that Recruits Smad2 to the TGFβ Receptor. Cell, 95(6), 779–791. 10.1016/S0092-8674(00)81701-8

Ueno, H., Huang, X., Tanaka, Y., & Hirokawa, N. (2011). KIF16B/Rab14 Molecular Motor Complex Is Critical for Early Embryonic Development by Transporting FGF Receptor. Developmental Cell, 20(1), 60–71. 10.1016/j.devcel.2010.11.008

van der Sluijs, P., Hull, M., Webster, P., Mâle, P., Goud, B., & Mellman, I. (1992). The small GTP-binding protein rab4 controls an early sorting event on the endocytic pathway. Cell, 70(5), 729–740. 10.1016/0092-8674(92)90307-X

Yap, C. C., & Winckler, B. (2012). Harnessing the power of the endosome to regulate neural development. Neuron, 74(3), 440–451. 10.1016/j.neuron.2012.04.015

Zhao, Y. G., & Zhang, H. (2019). Autophagosome maturation: An epic journey from the ER to lysosomes. Journal of Cell Biology, 218(3), 757–770. 10.1083/jcb.201810099

Zoghbi, H. Y., & Bear, M. F. (2012). Synaptic Dysfunction in Neurodevelopmental Disorders Associated with Autism and Intellectual Disabilities. Cold Spring Harbor Perspectives in Biology, 4(3), a009886–a009886. 10.1101/cshperspect.a009886

